# A cost-effective maize ear phenotyping platform enables rapid categorization and quantification of kernels

**DOI:** 10.1101/2020.07.12.199000

**Authors:** Cedar Warman, Christopher M. Sullivan, Justin Preece, Michaela E. Buchanan, Zuzana Vejlupkova, Pankaj Jaiswal, John E. Fowler

## Abstract

High-throughput phenotyping systems are powerful, dramatically changing our ability to document, measure, and detect biological phenomena. Here, we describe a cost-effective combination of a custom-built imaging platform and deep-learning-based computer vision pipeline. A minimal version of the maize ear scanner was built with low-cost and readily available parts. The scanner rotates a maize ear while a cellphone or digital camera captures a video of the surface of the ear. Videos are then digitally flattened into two-dimensional ear projections. Segregating GFP and anthocyanin kernel phenotype are clearly distinguishable in ear projections, and can be manually annotated using image analysis software. Increased throughput was attained by designing and implementing an automated kernel counting system using transfer learning and a deep learning object detection model. The computer vision model was able to rapidly assess over 390,000 kernels, identifying male-specific transmission defects across a wide range of GFP-marked mutant alleles. This includes a previously undescribed defect putatively associated with mutation of Zm00001d002824, a gene predicted to encode a vacuolar processing enzyme (VPE). We show that by using this system, the quantification of transmission data and other ear phenotypes can be accelerated and scaled to generate large datasets for robust analyses.

**One sentence summary:** A maize ear phenotyping system built from commonly available parts creates images of the surface of ears and identifies kernel phenotypes with a deep-learning-based computer vision pipeline.

## Introduction

High-throughput plant phenotyping is rapidly transforming crop improvement, disease management, and basic research (reviewed in (Fahlgren et al., 2015; Mahlein, 2016; Tardieu et al., 2017)). High-throughput phenotyping methods have been developed in several agricultural and model plant systems, including maize. There has been substantial progress towards deploying maize phenotyping systems, both in the private (Choudhury et al., 2016) and academic (Miller et al., 2017) realms. Many existing systems focus on phenotyping maize roots (Clark et al., 2013; Jiang et al., 2019) and above-ground shoots (Chaivivatrakul et al., 2014; Junker et al., 2014; Choudhury et al., 2016; Zhang et al., 2017). Maize ears, with the kernels they carry, contain information about the plant and its progeny. Ears are easily stored, and do not require phenotyping equipment to be in place in the field or greenhouse at specific times during the growing season. Ears are a primary agricultural product of maize, which has led the majority of previous phenotyping efforts to focus on aspects of yield, such as ear size, kernel row number, and kernel structure and dimensions (Liang et al., 2016; Miller et al., 2017; Makanza et al., 2018). These studies have used techniques that varied from expensive and specialized three-dimensional or line-scanning cameras (Liang et al., 2016; Wen et al., 2019) to relatively low-cost flatbed scanners and digital cameras (Miller et al., 2017; Makanza et al., 2018).

Beyond their agricultural importance, studying maize ears can answer fundamental questions about basic biology. The transmission of mutant genes can be easily tracked in maize kernels by taking advantage of a wide variety of visible endosperm markers (Neuffer et al., 1997; Li et al., 2013), which can be genetically linked to a mutant of interest (e.g. (Arthur et al., 2003; Phillips and Evans, 2011; Bai et al., 2016; Huang et al., 2017; Warman et al., 2020)). On the ear, kernels occur as an ordered array of progeny, which allows the transmission of mutant alleles to be tracked not only by total transmission for each individual cross, but within individual ears. Historically, transmission of markers has been quantified by manual counting. This approach has several limitations, among them a lack of a permanent record of the surface arrangement of kernels on the ear. The same disadvantages apply to most high-throughput kernel phenotyping methods, which generally rely on kernels being removed from the ear before scanning and do not typically include marker information.

Computer vision approaches to automated kernel counting can improve throughput in phenotyping systems and improve the quality of data collected by including positional information for each kernel. One central challenge is successfully identifying which parts of an image contain the objects of interest and which parts contain the background, either through object detection (drawing a bounding box around the object) or segmentation (assigning each pixel in the image as “object” or “not object”). Previous systems have taken advantage of plant color or edges to algorithmically separate objects for quantification in some specialized contexts (Zhang et al., 2017; Makanza et al., 2018). These approaches can be computationally efficient, but are limited by variations in lighting conditions, image quality, and the distribution of objects in an image. Closely packed objects, such as kernels on a maize ear, can be difficult to separate using these methods, especially when the objects do not have consistent colors or clear edges.

Some of these obstacles to object detection can be overcome with deep learning approaches. These approaches have been applied to a variety of biological problems, and can show dramatic improvements over traditional methods (reviewed in (Angermueller et al., 2016; Ching et al., 2018)). Deep learning uses the fundamental concept of artificial neural networks, in which multiple nodes (sometimes referred to as neurons) are arranged in variously connected layers. Nodes have associated parameters that are adjusted as the model is exposed to data.

Data moves through the network from an input layer to at least one hidden layer, and finally to the output layer. Deep learning is characterized by a neural network with multiple hidden layers, in which each layer describes features of the data being passed through the network (Ching et al., 2018). A subset of deep learning approaches called convolutional neural networks (CNNs) are particularly useful for image analysis. CNNs contain at least one convolutional layer, in which a filter moves (convolves) across an image to abstract information into the layer (Rawat and Wang, 2017). CNNs form the foundation of the object detection methods implemented in Tensorflow (e.g. Object Detection API, (Huang et al., 2016)) and Darknet (e.g. YOLO (Redmon and Farhadi, 2018)) that have seen widespread use across disciplines. Examples of such networks being used in biological contexts include plant disease detection (Mohanty et al., 2016), leaf quantification (Ubbens and Stavness, 2017), inflorescence movement tracking (Gibbs et al., 2019), and hypocotyl segmentation (Dobos et al., 2019).

Here we describe a novel maize ear phenotyping system and computer vision pipeline. The maize ear scanner and image processing pipeline is a cost-effective method to improve ear phenotyping. The design described here is built from easily acquired parts and a basic camera, making this approach accessible to most if not all labs. Using the maize ear scanner, flat projections of roughly cylindrical maize ears can be produced that provide a digital record of the surface of the ear. These projections can then be quantified in a variety of ways to track the locations and identities of kernel phenotypes, including marker genes. In addition, projections can be quantified for kernel phenotypes and locations with a deep-learning-based computer vision pipeline implemented in Tensorflow, a free and open source framework (Abadi et al., 2016a). Finally, we use the system to analyze a large dataset of ears to assess mutant effects on transmission rate. We demonstrate that this system substantially increases phenotyping throughput, enabling more rapid biological discovery and more thorough quantitative analyses.

## Results

### Design and construction of the maize ear scanner

We designed a simple, custom-built scanner to efficiently phenotype maize ears (Maize Ear Scanner, MES). The MES rotates each ear 360° while a stationary camera records a video, which can then be processed into a cylindrical projection. Materials for constructing the scanner were limited to those that are widely available and affordable (Table 1). The frame of the scanner was built from dimensional lumber, with a movable mechanism built from drawer slides that enables a wide range of ear sizes to be accommodated (Figure 1A). A rotisserie motor spins the ear at a constant speed while a USB camera or cell phone camera records a video of the rotating ear. The scanning process, including the insertion of the ear into the scanner and video capture, takes approximately 1 minute per ear.

**Figure 1.**
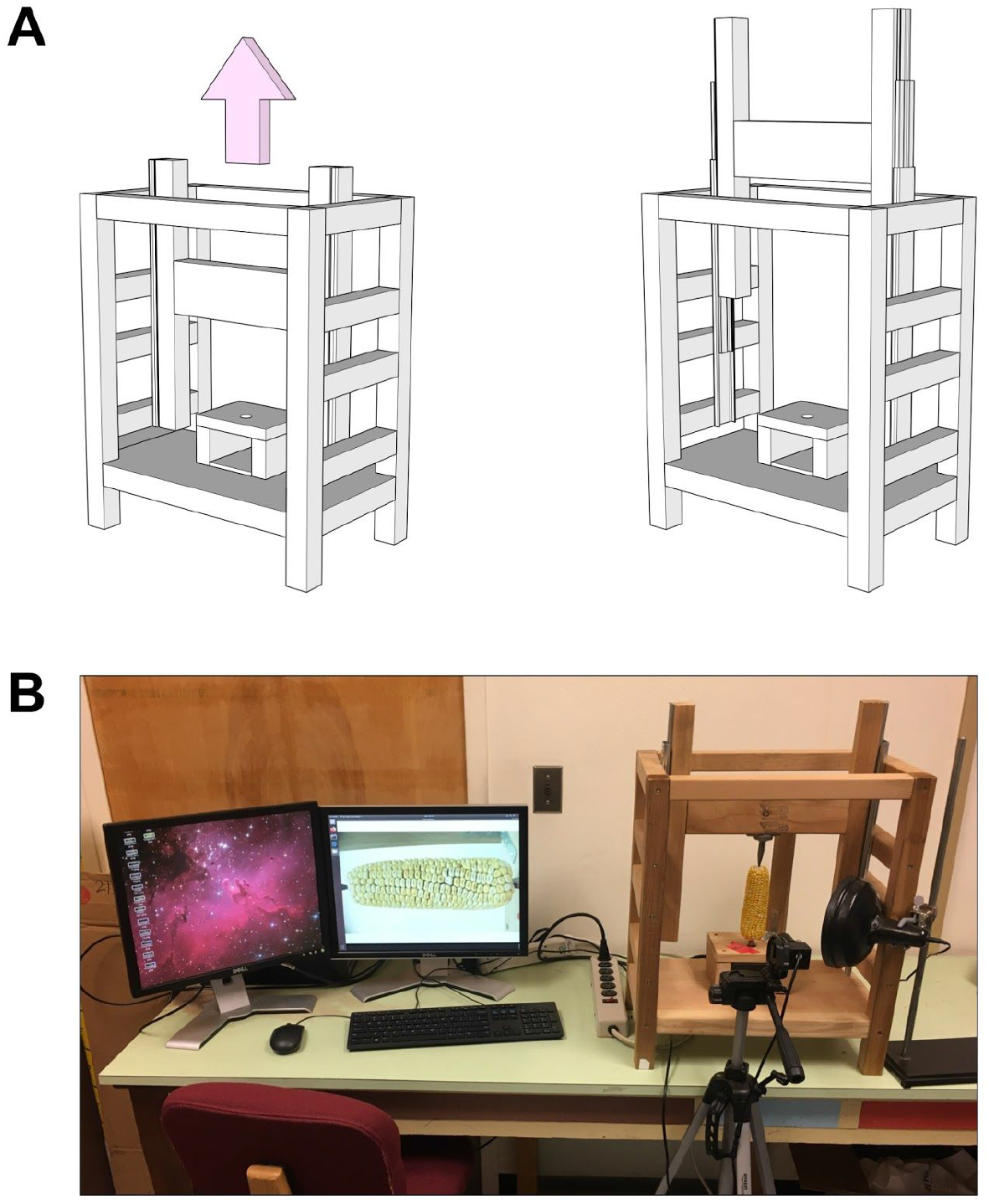
Efficient, cost-effective maize ear phenotyping with rotational scanner. **(A)** Schematics of maize ear scanner in closed position (left) and open position (right). Full construction diagrams are available in Supplemental File 1. **(B)** Image of scanner with ear in place. A dedicated USB camera is positioned in front of the ear as shown, with the ear centered in the frame. A video is captured as the ear spins through one full rotation, which is then processed to project the surface of the ear onto a single flat image. An optional blue light source for GFP imaging is shown on the right side of the scanner.

**Table 1.**
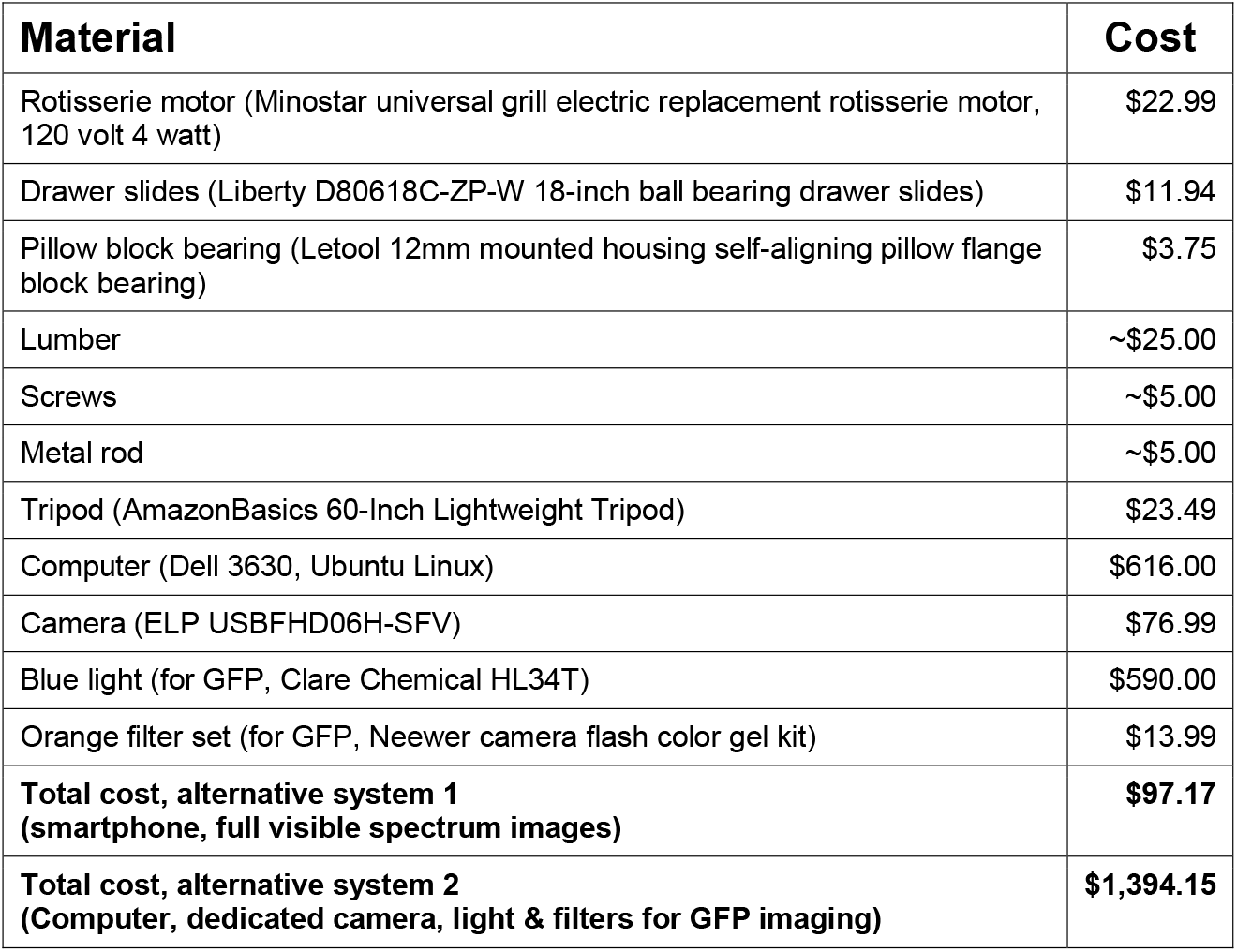
Materials and costs for scanner construction

We tested two configurations of the scanning system. In the first, a minimal configuration, a cell phone camera was used to capture movies of the rotating ear in full spectrum visible light (MES v1.0). This configuration cost less than $100 (Table 1), excluding the cost of the cell phone, and is capable of producing flat projections from a variety of ears in visible light. The second configuration uses a dedicated USB camera driven by a computer (MES v2.0, Figure 1B). This configuration costs about $1,400 (Table 1), including a blue light and orange camera filter to capture GFP kernel markers present in a population of transgenic mutants (Li et al., 2013). The second configuration increases the scanner’s efficiency by automating video processing, annotation, and distribution to cloud or local storage systems.

### Processing videos into projections for manual quantification

The output of the scanner is a video of the rotating ear. This video can be directly quantified, but we found a ‘flat’ image projection most useful for visualizing the entire surface of the ear, as well as for quantifying the distribution of kernel phenotypes. To produce this projection with videos captured by an external camera or cell phone, videos were first uploaded to a local computer and annotated with identifying metadata. This process was streamlined in the second configuration of the scanner. In this configuration, videos were captured directly to the computer using the command line utility FFmpeg (version 3.4.6) to control a USB camera. Videos were automatically processed each night, with the resulting projections uploaded to cloud storage (Figure 2A).

**Figure 2.**
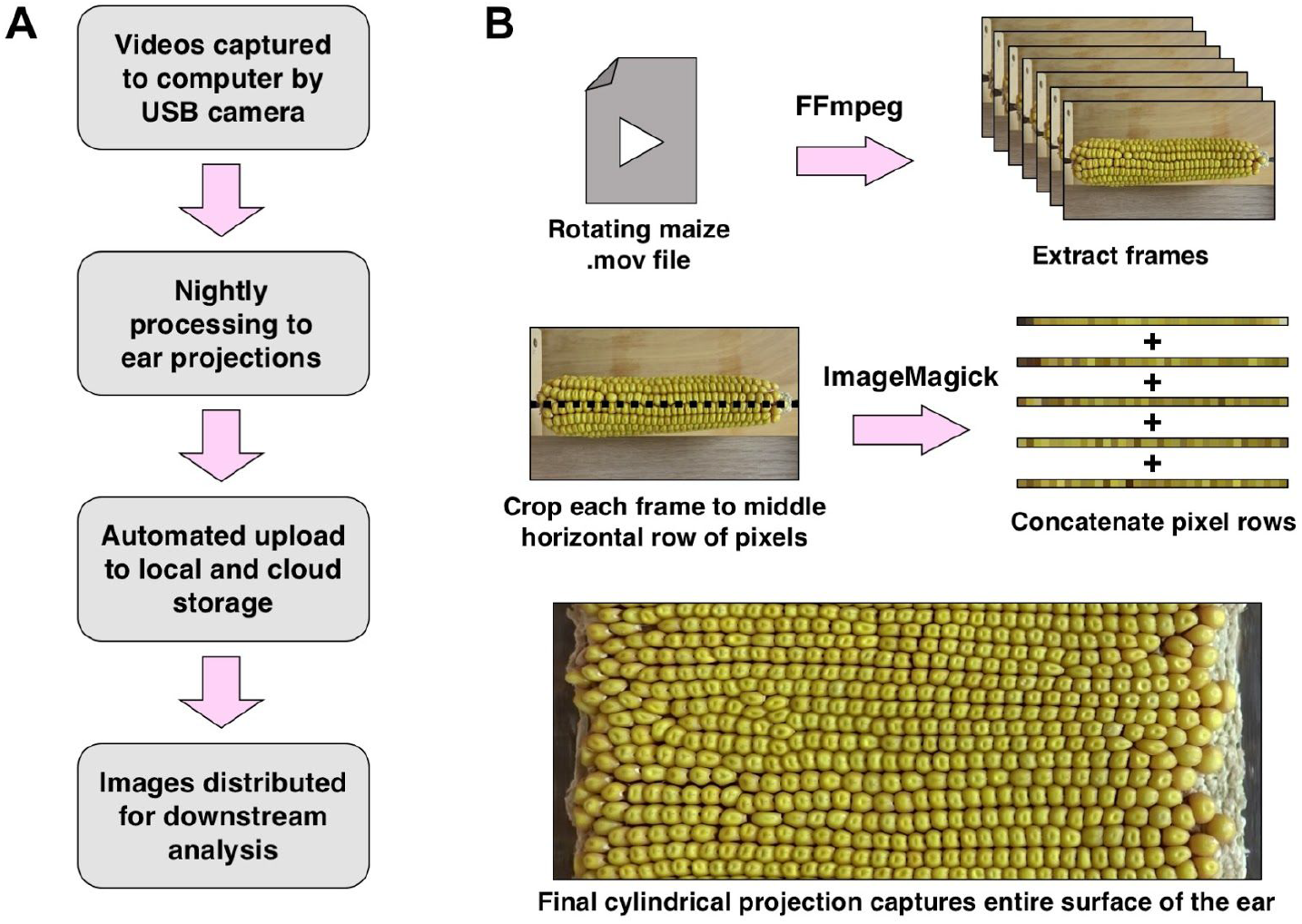
Processing videos into flat ear projections. **(A)** Video annotation and processing workflow. **(B)** Processing videos to flat ear projections. The process of generating a projection from a video begins by extracting individual frames using FFmpeg. After frames are extracted, each frame is cropped to the middle horizontal row of pixels using the command line utility ImageMagick. The resulting collection of pixel rows, one per frame, is then concatenated into a single image depicting the entire surface of the ear.

Video processing consists of three steps (Figure 2B). In the first, FFmpeg is used to extract frames from the video into separate images. Next, images are cropped to the center horizontal row of pixels using the command line utility ImageMagick (version 6.9.7). Finally, all rows of pixels, one from each frame, are appended sequentially, resulting in the final image. Due to the scanner’s consistent rotational speed, a fixed number of frames cover one complete rotation, resulting in no gaps or overlap in ear projections.

Images of a variety of maize ears representing several widely used kernel markers were captured using the scanner (Figure 3A). Both anthocyanin (*c1, a2*, and *pr1*) and fluorescent (*Ds-GFP*) kernel markers were clearly discernible in the final images, as well as the kernel morphology marker *brittle endosperm1* (*bt1*). Digital projections were manually quantified for color and fluorescent kernel phenotypes using the FIJI distribution of ImageJ (Figure 3B). Using this approach, annotation of an entire ear could be completed in 5 to 10 minutes, depending on the size of the ear and the relative experience level of the annotator. In addition to producing total quantities of each kernel phenotype, manual annotations result in coordinates for each annotated kernel, which can be further analyzed if desired. Manual annotations of scanner images in ImageJ were compared to manually counting the kernels on the ear (Figure 3C). We observed a significant correlation between these two methods (R^2^ > 0.999), validating the scanning method. To test the utility of the maize ear scanner, we scanned and manually counted over 400 ears with marker-linked mutations in >50 genes. With these methods, we were able to detect weak but significant transmission defects (~45% transmission of a marker-linked mutation) for a number of mutant alleles, using both anthocyanin and GFP kernel markers. Manually counted ear scanner image validation is described in detail in a previous study (Warman et al., 2020).

**Figure 3.**
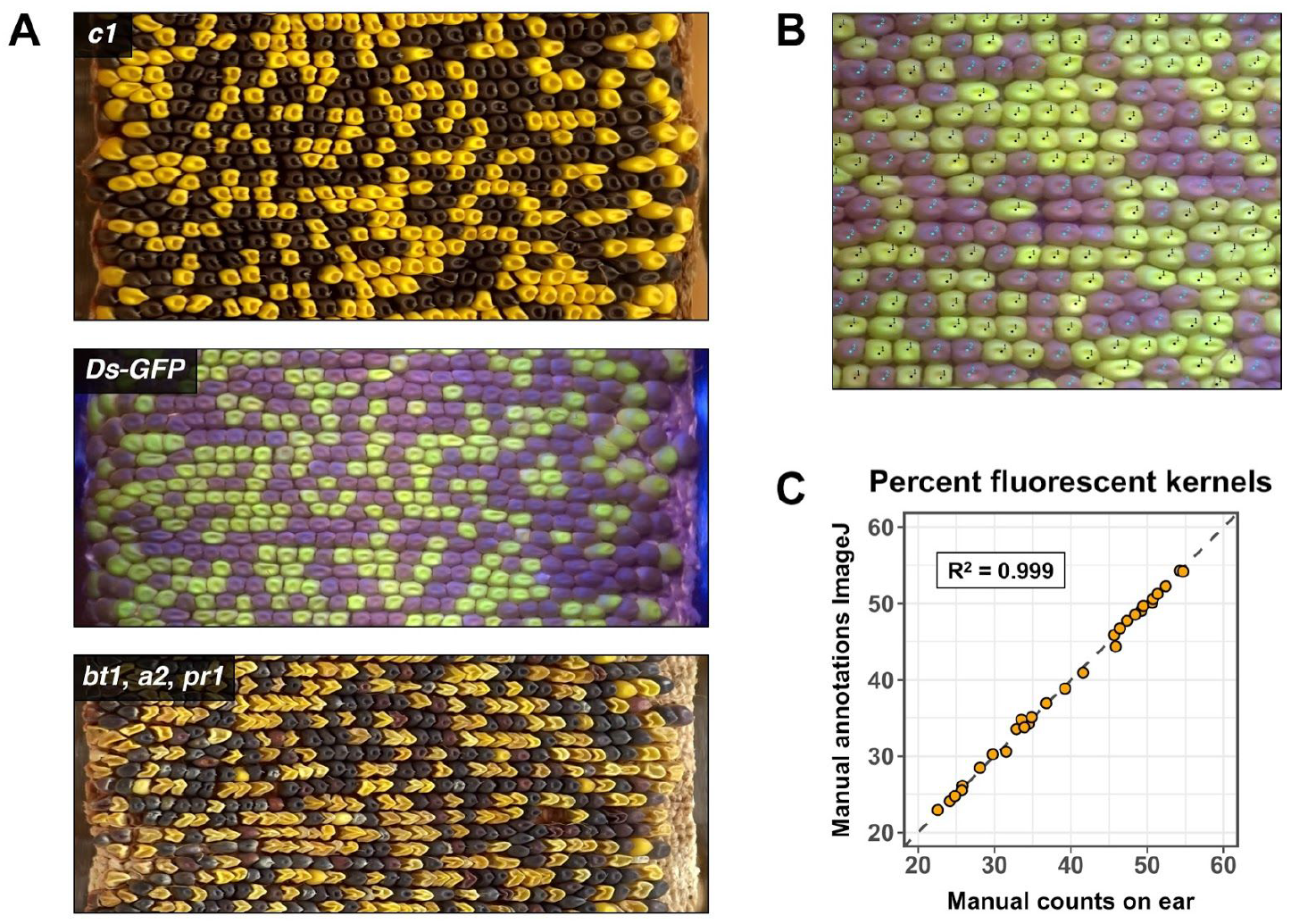
Examples of ear surface projections and manual quantification using ImageJ. **(A)** Representative ear projections for several widely-used maize kernel markers. From top to bottom: anthocyanin gene *c1*; GFP fluorescent kernel marker *Ds-GFP;* anthocyanin and kernel morphology markers *bt1, a2*, and *pr1*. **(B)** Representative example of manual annotation of fluorescent maize kernels. An ear of maize segregating for fluorescence was imaged. Fluorescent (black dots) and non-fluorescent (blue dots) kernels were manually identified using the ImageJ Cell Counter plugin. **(C)** Comparison of manually counting kernels on ears vs manually annotating kernels from ear projections using ImageJ. Fluorescent and non-fluorescent kernels were counted, with the percentage of kernels showing fluorescence shown here.

### A traditional computer vision approach for automated discrimination of fluorescent and wild-type kernels

To increase throughput, we investigated computer vision methods to identify kernel locations and phenotypes from two-dimensional ear projections. We were most interested to establish a pipeline to automate the counting of *Ds-GFP* vs. wild-type (non-fluorescent) kernels, due to the broadly applicable use of this marker to quantify transmission rates for several thousand available mutations (Li et al., 2013; Warman et al., 2020). First, a traditional computer vision approach was assessed for its feasibility for quantification of images with GFP kernel markers. In this method, region-based segmentation of a two-dimensional ear projection was performed using a watershed transformation followed by morphological opening to segment individual kernels (Supplemental Figure 1). We found that extracting the blue channel of the RGB image for segmentation avoided inaccuracies resulting from varying intensities of kernel fluorescence in the green and red channels. After segmentation, segments were classified using k-means clustering into two groups for presence and absence of GFP. Fine-tuning of watershed parameters resulted in the accurate segmentation of individual images (Supplemental Figure 1A). However, because of variations in lighting, GFP intensity, kernel shape, and spacing on the ear, this method generalized poorly across a larger test dataset (Supplemental Figure 1B). This method was able to predict total fluorescent and non-fluorescent kernel numbers with some success (linear regression, adjusted R^2^ = 0.186, 0.205, respectively), but failed to accurately predict the percentage of kernels carrying the GFP marker (linear regression, adjusted R^2^ = 0.000). Because marker-tagged mutants show Mendelian (50%) or near-Mendelian inheritance, accurate counts are required to measure abnormal inheritance with sufficient statistical significance.

### Implementation of a deep learning model for automated kernel detection

To overcome the large variation in ear images, we turned to deep learning models, which are effective in detecting objects within heterogeneous images. Models using convolutional neural network architecture (CNNs) have dominated performance metrics in the computer vision field for several years (for example, Collection Of Common Objects (COCO) Object Detection Challenge, http://cocodataset.org/#detection-leaderboard, Open Images Object Detection Challenge, https://storage.googleapis.com/openimages/web/challenge2019.html#object_detection). We used the TensorFlow library (Abadi et al., 2016b) and Object Detection API (Huang et al., 2016) to implement a CNN-based model for our purposes. A tradeoff between speed and accuracy exists when implementing deep learning models for object detection. Some models, such as YOLO (Redmon and Farhadi, 2018) are significantly faster than the models available in the TensorFlow Object Detection API. However, because the image processing could be run independently of ear scanning, even several minutes of processing time per image would not present a bottleneck. For the pipeline, we chose to use the Faster R-CNN with Inception Resnet v2, with Atrous convolutions (Ren et al., 2015; Szegedy et al., 2016). This model was selected to balance speed and accuracy for our application based on its performance on the COCO dataset. (https://github.com/tensorflow/models/blob/master/research/object_detection/g3doc/detection_model_zoo.md).

CNNs require training data to generate effective models. To train the network, we generated a dataset of 300 scanned ear images with all kernels annotated with bounding boxes and marker classes, either fluorescent or non-fluorescent. Images were generated by scanning ears produced from heterozygous outcrosses of mutant alleles tagged with GFP fluorescent kernel markers, with 150 scanned ear images for each field season. The mean kernel number for training ear images was 349, resulting in >100,000 bounding boxes in the training set. We used a transfer learning approach because of the large amount of training data required to accurately train a neural network from scratch. Transfer learning takes advantage of a well-trained network (in this case trained on the COCO dataset, >200,000 images with objects in 80 categories labeled with bounding boxes) to form the foundation for a new network optimized for a specific task. The weights of the inner layers of the network are updated based on the new training data, and the output layer is modified to reflect the new classes (fluorescent and non-fluorescent).

Our first attempts at training the network led to poor results (Figure 4A). Kernel bounding boxes were accurate in the top portion of test images, but these results failed to generalize across the entire image. Due to the large number of kernels on each image (over 600 on some ears), we suspected graphics processing unit (GPU) memory limitations may have caused incomplete annotations. Supporting this explanation, we gained incremental improvements by running the training and testing on a GPU with more memory (Nvidia V100 with 32 GB of memory versus an Nvidia M10 with 8 GB of memory) and a configuration that increased the number of initial bounding box proposals in the model.

**Figure 4.**
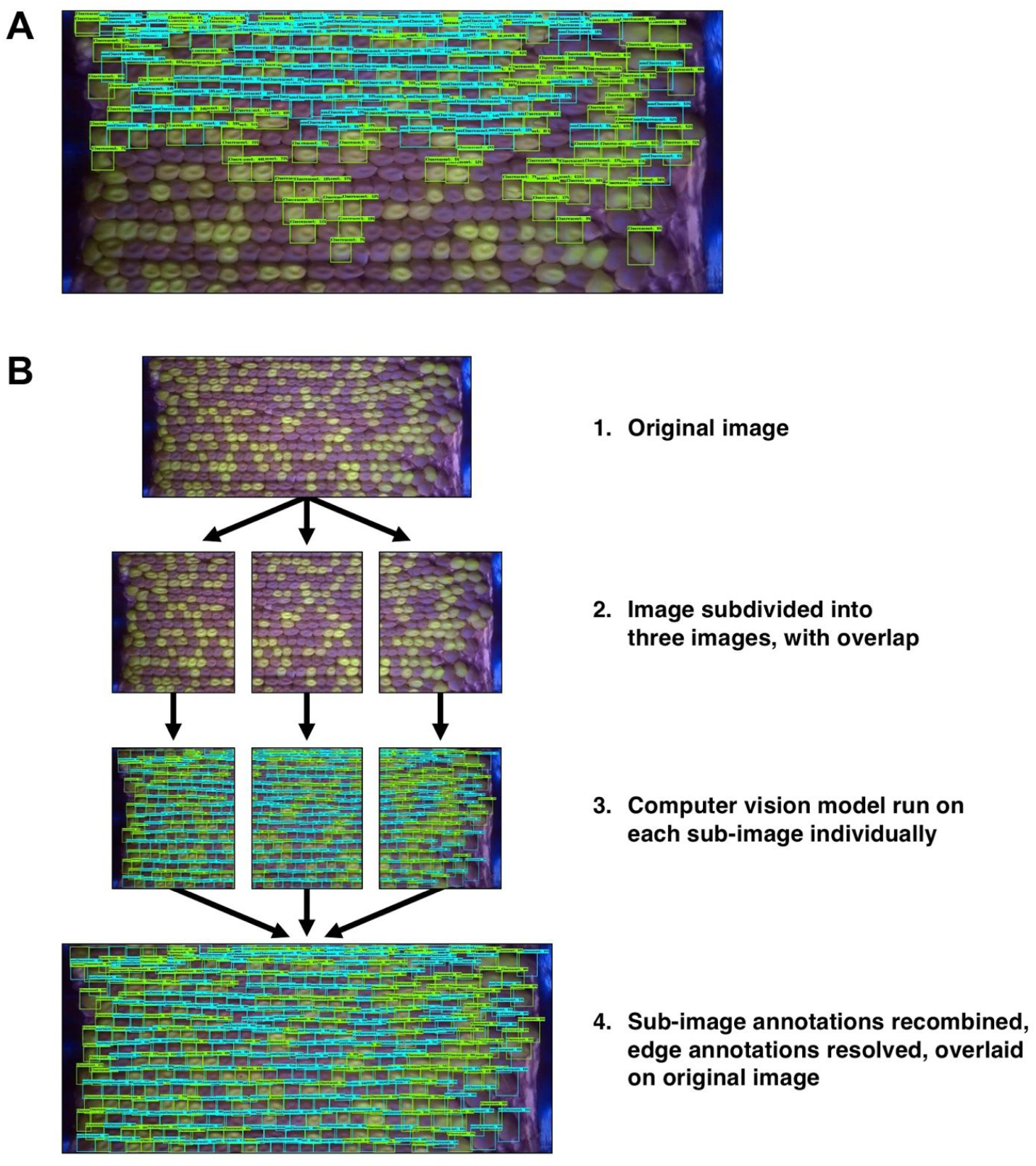
Workflow for subdividing images during model training and inference. **(A)** Representative image of initial object detection attempts showing incomplete bounding box annotations. Annotations were biased towards the top of the image, and failed to identify the majority of kernels. **(B)** Image subdivision workflow. Images were first subdivided into three smaller images, with overlap between the images. The computer vision model was then run on each sub-image individually. Bounding boxes near the vertical borders between sub-images were removed to avoid split bounding boxes on single kernels. Annotations were recombined, then redundant boxes in overlapping sections were removed with non-maximum suppression. Finally, completed annotations were overlaid on the original image.

One way to reduce the computational power necessary for a deep learning task is to subdivide the task into a series of simpler problems. In this case, we chose to subdivide each image into three sub-images, both for the training and for the testing of the model (Figure 4B). Images were subdivided vertically, with overlapping regions included between each division. After images were subdivided, the model was run on each sub-image individually. Bounding boxes near the vertical divisions were then removed to avoid partial bounding boxes for kernels that spanned two sub-images. Finally, annotations for the three sub-images were combined, and redundant bounding boxes in the overlapping areas were removed with non-maximum suppression, a process that resolves redundant bounding boxes by comparing their overlap and confidence scores. This method reduced the GPU memory required for inference, and resulted in accurate annotations across entire images.

### Deep learning models trained on images from individual cameras improved detection of kernels and phenotypic classes

To test the deep learning models, we created a dataset of scanned ear images from the 2018 and 2019 field seasons, with 160 images from each season. Ears were generated from reciprocal outcrosses (heterozygous mutants crossed to wild-type lines both through the male and female), and were manually annotated with ImageJ to produce total fluorescent and non-fluorescent kernel counts for each ear. Testing set ear images were not used for training or validation of the model. Lines represented mutant alleles in a variety of genes highly expressed in the maize male gametophyte (Warman et al., 2020). Kernels containing mutant alleles were marked with a GFP seed marker originating from *Ds-GFP* transposable element insertions. Projections generated across the two seasons represented a wide range of ears. Variations found in projections included differences in kernel size, shape, GFP intensity, and color. In addition, different cameras were used in each year, representing MES v1.0 and v2.0.

We first aimed to create a model with as much generalizability as possible, and thus included training images from both years. This first model, trained on images from two cameras, detected kernels in a test dataset with a moderate degree of accuracy (Supplemental Figure 2). We used adjusted R^2^ values as the principal performance metric, comparing total fluorescent and non-fluorescent kernel counts between model predictions and manual annotations. The closer the R^2^ value is to one, the closer the model predictions are to manual counts, indicating higher accuracy. The resulting percentage of fluorescence kernels was quantified with an adjusted R^2^ of 0.930. In addition, we calculated the mean absolute deviation in kernel count across the entire test dataset. The mean absolute deviation for fluorescent kernels was 5.87, whereas the mean absolute deviation for non-fluorescent kernels was 11.92. The mean absolute deviation for percent fluorescent kernel transmission was 1.85%.

A single model trained on a combined dataset from both years accurately identified kernels in scanned ear images. However, training separate models for each year substantially increased overall performance across a wide variety of images from both years of our test dataset (Figure 5). Individual models were robust to variations in kernel appearance, as well as to variations in ear size and kernel spacing. The models predicted total fluorescent and non-fluorescent kernels across the 2018 and 2019 test datasets with a high degree of accuracy (Figure 5B-C). The resulting transmission rate predictions were accurate across a wide range of inheritance values for both years (linear regression; adjusted R^2^ = 0.984, 0.945, respectively). The mean absolute deviations for fluorescent kernels in individual models for 2018 and 2019 were 5.74 and 5.75, respectively, whereas the mean absolute deviations for non-fluorescent kernels were 8.58 and 6.81. The mean absolute deviations for percent fluorescent kernel transmission were 0.885% and 1.38%. Training individual models was substantially faster than training a single model (~100-fold faster training time on an Nvidia V100 GPU). Detailed metrics for model training can be found in the Materials and Methods section below. While the variation introduced by using different cameras for each year was likely responsible for the increased accuracy of individual models, we cannot rule out other potentially correlated factors in the two growing seasons. Because of their increased accuracy, we proceeded to use individual models for each camera/year to investigate transmission rates for *Ds-GFP* mutant alleles. We term these models collectively as EarVision.

**Figure 5.**
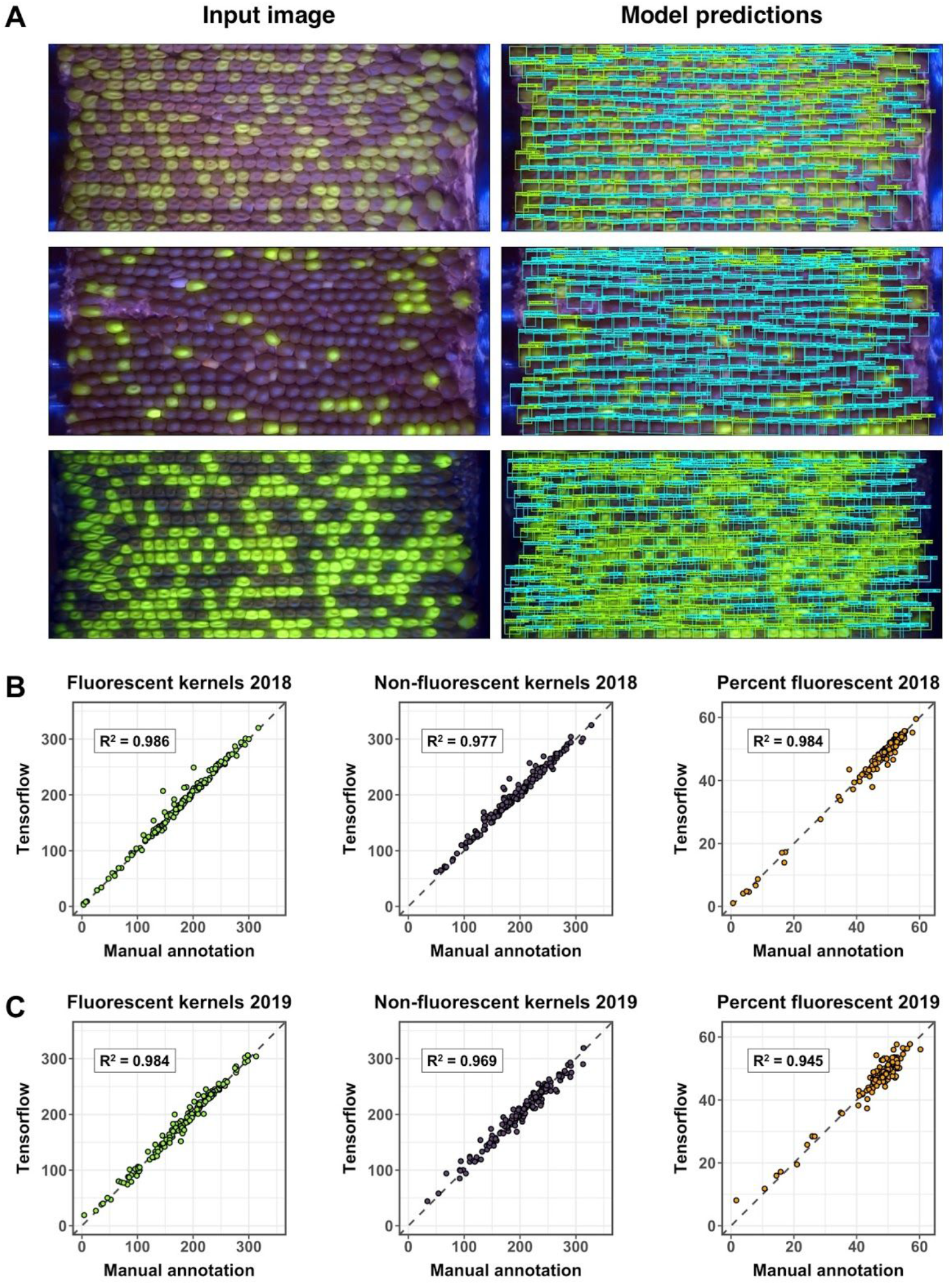
Deep learning models trained on image datasets from different field seasons and cameras accurately detected kernels and classes across a test dataset. **(A)** Example test images and annotations predicted by model. Top images: A typical ear from the 2018 field season showing Mendelian inheritance of GFP-marked kernels. The model predicted a 45% transmission rate, whereas manual annotation indicated 46.3% transmission. Middle images: a 2018 ear showing significant transmission defect, with few GFP-marked kernels. The model predicted a 16.9% transmission rate, whereas manual annotation indicated 16.2% transmission. Bottom images: 2019 ear showing Mendelian inheritance. The model predicted a 50.1% transmission rate, whereas manual annotation indicated 50.7% transmission. **(B)** Total kernel counts and percent GFP across the 2018 test dataset (160 images). Adjusted R^2^ values were calculated using a linear model comparing manual counts (x-axis) to deep learning model predictions (y-axis). Dashed diagonal lines represent equal values for both manual counts and model predictions. Adjusted R^2^ values for total fluorescent, non-fluorescent, and percent fluorescent kernels were above 0.97. **(C)** Total kernel counts and percent GFP across the 2019 test dataset (160 images). Adjusted R^2^ values were again calculated using a linear model comparing manual counts to deep model predictions, and were above 0.94.

### Application of deep learning models to a large ear projection dataset

To test the EarVision deep learning models on a larger dataset, we quantified a set of 369 scanned ear images that had manually counted kernels from a previous study (Warman et al., 2020). The original dataset consisted of images of ears from maize plants grown during the 2018 field season. Ears were harvested from different plants with single *Ds-GFP* insertions in 44 genes. A total of 48 mutant alleles were examined, with four genes having two independent *Ds-GFP* insertions. Genes were selected because they are highly expressed in the male gametophyte. Reciprocal outcrosses of heterozygous mutants were carried out to functionally interrogate these genes. This process led to the identification of several mutant alleles with reduced transmission through the male. We assessed the accuracy of the EarVision model’s predictions by comparing transmission rates for manually annotated images and model predictions (Supplemental Figure 3). For crosses through the female, the model predicted that mutations in all 44 genes had no significant difference from Mendelian (50%) inheritance, consistent with manual annotations (Supplemental Figure 3A). For crosses through the male, the model successfully predicted 7/8 alleles that showed significant transmission defects when transmission was quantified manually, with the transmission of one of these alleles predicted as nonsignificant by the model (Supplemental Figure 3B). A generalized linear model showed no evidence of significant systematic differences between manual annotations and model predictions (p-value > 0.8).

A second set of reciprocal crosses was carried out in the 2019 field season to increase the size of the ear image dataset. Crosses from the 2019 field season included plants with previously tested mutant alleles to determine whether transmission rates identified in 2018 remained consistent in the following year. Crosses also contained plants with additional alleles that were not included in the published analysis (Warman et al., 2020), either because of insufficient crosses in 2018 (5 alleles) or lack of PCR confirmation of *Ds* insertion location (7 alleles). In total, approximately 1000 ears from plants containing 60 mutant alleles were quantified using individual computer vision models for 2018 and 2019 field seasons. Combined 2018+2019 model estimates were largely aligned with 2018 manual annotations for both male and female crosses (Figure 6). The data from the combined models correctly predicted no significant transmission defects through the female for 56/60 alleles in the combined dataset, with 4/60 alleles assigned GFP transmission rates significantly increased over Mendelian inheritance (Figure 6A). These apparent false positives were likely the result of a systematic undercount of non-fluorescent kernels in a small subset of female crosses in the 2019 dataset (Supplemental Figure 4A-B). This is potentially due to the relatively strong GFP signal arising from doubled dosage of *Ds-GFP* in the endosperm, leading to reduced accuracy in the recognition of non-fluorescent kernels (Supplemental Figure 4C). The model correctly predicted all 8 alleles showing significant transmission defects as determined by 2018 manual counts (Figure 6B). In addition, the model predicted male transmission rates for the 12 alleles not present in the 2018 dataset, the majority of which (11/12) showed no evidence of non-Mendelian inheritance. However, the model identified a significant, previously undescribed, male-specific transmission defect associated with a *Ds* insertion predicted to be in the maize gene Zm00001d002824 (Table 2). Zm00001d002824 codes for a putative vacuolar-processing enzyme (VPE). VPEs have been shown to be involved in the maturation of vacuolar proteins as well as vacuolarūorganized programmed cell death (Yamada et al., 2005), and their potential role in male gametophyte function is unexplored.

**Figure 6.**
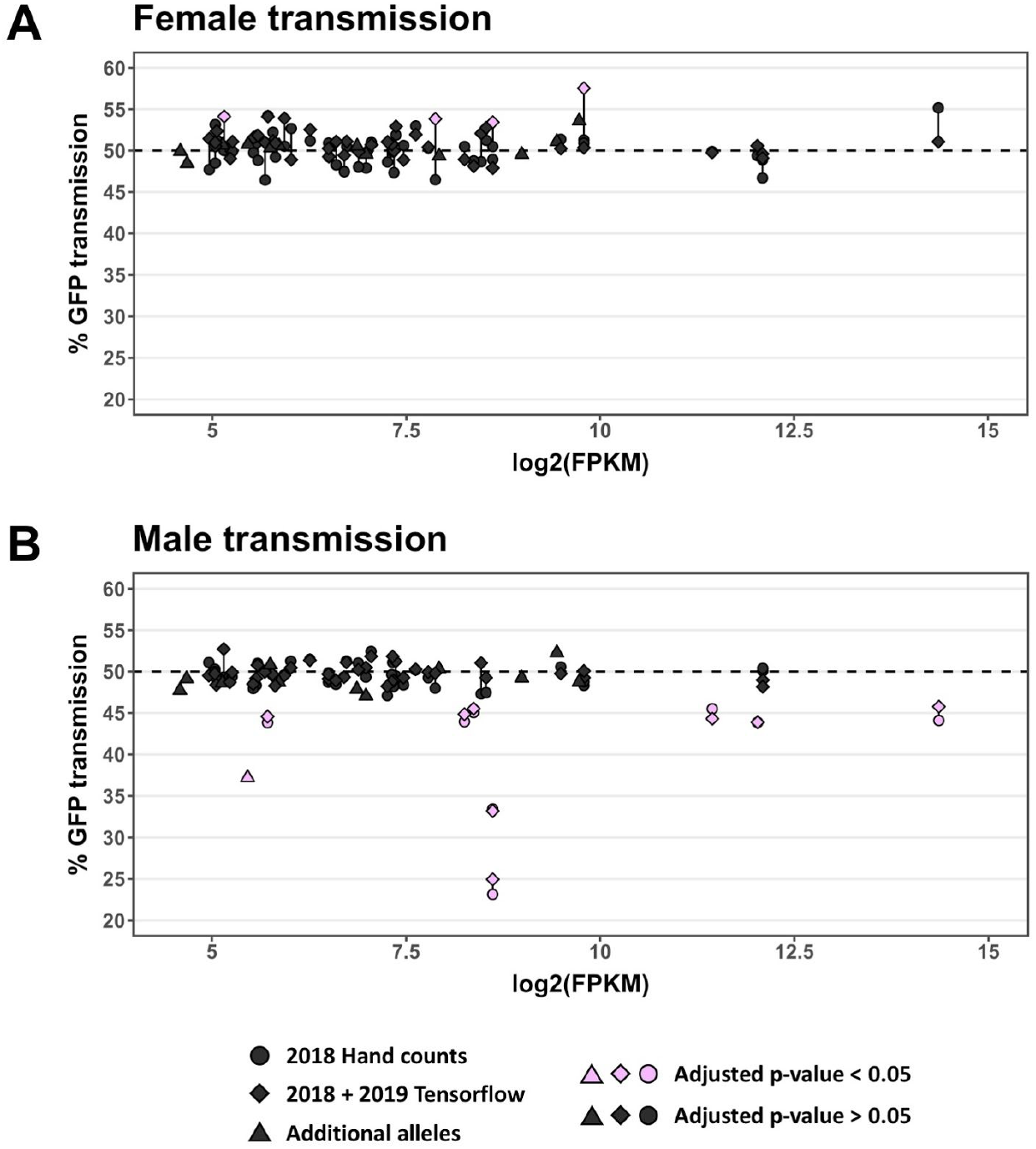
Computer vision model predictions align with manual counts for GFP transmission across two field seasons. **(A)** Transmission results from plants containing heterozygous *Ds-GFP* insertion alleles outcrossed through the female. A total of 60 alleles were quantified in individual plants. Genes with mutant alleles belonged to three categories (Seedling Only, Vegetative Cell, Sperm Cell) based on high expression levels in those tissues, shown on the x-axis in log2(FPKM). **(B)** Transmission rates for all alleles assessed, when plants containing heterozygous *Ds-GFP* insertion alleles were outcrossed through the male. Expression levels in previously described categories are shown on the x-axis.

**Table 2.** Characteristics of genes harboring newly-assessed *Ds-GFP* alleles

| Expression Category | Reason for exclusion from Warman et al. 2020 | Gene designation (v3) | Gene designation (v4) | Ds-GFP allele | Male transmission rate | Adjusted p-value | Best BLAST Hit, *A. thaliana* | Annotation (B73v4 Gramene) |
| --- | --- | --- | --- | --- | --- | --- | --- | --- |
| seedling_only | Insufficient crosses | GRMZM2G080724 | Zm00001d031325 | tdsgR106E07 | 48.64% | 0.589 | AT4G27670 | Heat shock protein 21 |
| seedling_only | Insufficient crosses | GRMZM2G148333 | Zm00001d005798 | tdsgR44E07 | 47.87% | 0.503 | AT3G14230 | Ethylene-responsive transcription factor RAP2-2 |
| seedling_only | Insufficient crosses | GRMZM2G148387 | Zm00001d017240 | tdsgR91G06 | 50.80% | 0.686 | AT5G63030 | Glutaredoxin-C1 |
| seedling_only | Insufficient crosses | GRMZM2G374302 | Zm00001d051194 | tdsgR65A10 | 52.26% | 0.258 | AT2G16500 | Arginine decarboxylase |
| sperm_cell_high | Insufficient crosses | AC194405.3_FG021 | Zm00001d012575 | tdsgR83A02 | 49.21% | 0.796 | AT1G19100 | Protein MICRORCHIDIA 6 |
| sperm_cell_high | unconfirmed insertion | GRMZM2G038851 | Zm00001d002570 | tdsgR06C04 | 48.76% | 0.716 | AT3G57870 | SUMO-conjugating enzyme SCE1 |
| sperm_cell_high | unconfirmed insertion | GRMZM2G062554 | Zm00001d002824 | tdsgR89B08 | 37.21% | <0.000001 | AT1G62710 | Vacuolar processing enzyme, beta-isozyme |
| sperm_cell_high | unconfirmed insertion | GRMZM2G124365 | Zm00001d012674 | tdsgR29A11 | 49.13% | 0.796 | AT3G29200 | Chorismate mutase1 |
| vegetative_cell_high | unconfirmed insertion | GRMZM2G033828 | Zm00001d031678 | tdsgR67H12 | 48.73% | 0.703 | AT3G12280 | Retinoblastoma family3 |
| vegetative_cell_high | unconfirmed insertion | GRMZM2G051491 | Zm00001d005053 | tdsgR85A08 | 50.32% | 0.890 | AT3G10870 | Methylesterase 17 |
| vegetative_cell_high | unconfirmed insertion | GRMZM2G140107 | Zm00001d042353 | tdsgR02A05 | 47.43% | 0.283 | AT1G04920 | Sucrose phosphate synthase2 |
| vegetative_cell_high | unconfirmed insertion | GRMZM2G319167 | Zm00001d039693 | tdsgR108A02 | 47.05% | 0.243 | AT3G27670 | Protein RST1 |

## Discussion

Large amounts of information can be obtained from maize ears. Certain types of information, such as yield and kernel quality, have direct relevance for improving maize for agricultural purposes. Other types of information, such as mutant transmission, can be used to study fundamental biological processes. Our goal was to develop a methodology to capture some of this information via digital imaging and automated kernel detection. This methodology enables standardized, replicable measurement of kernel phenotype distribution, as well as provides a permanent digital record of ears for archiving and future reanalyses. The scanner is fast and cost-effective in its minimal configuration (MES v1.0). A step-up from the minimal configuration (MES v2.0) enables a more automated system for file transfer and video-to-projection generation, and the addition of EarVision, enabling deep-learning-based kernel quantification from the resulting images, dramatically scales up the amount of quantitative data that can be feasibly generated. In addition, automated quantification avoids variation introduced by multiple individuals manually quantifying images.

While the scanner provides useful phenotyping data, it has some notable limitations. Cylindrical projections are a convenient way of visualizing the entire surface of an ear in a single image. However, because maize ears are not perfect cylinders, the projections distort regions of the ear that are not cylindrical, typically the top and bottom, resulting in kernels that appear larger than those in the middle of the ear (Figure 3A). Excessively curved ears, sometimes resulting from uneven pollination, can also lead to severe distortions. Because of these distortions, measuring qualities like kernel and ear dimensions can be challenging. While approximate values for these metrics can be calculated, in the future more precise measurements could potentially use the source video as input to model the ear in three dimensions, particularly with the addition of calibration objects (Feldmann et al., 2019). Thus, we have limited the use of the EarVision pipeline to relatively straight, uniform-thickness ears, whereas more distorted ear images can be quantified following manual annotation.

Capturing high-quality and standardized images is crucial for the system. Differences in photography equipment, image quality, and variation in ear and kernel morphology can compromise accuracy. A small subset of kernels that were significantly outside the normal range of color variation were not identified by the model, particularly in images with non-optimal exposure (Supplemental Figure 4). These cases can likely be resolved with an improved imaging protocol and quality control, particularly of high contrast, strong GFP signal images. The overall impact of these weaknesses on the model’s accuracy was low, due to their relative scarcity.

The EarVision pipeline appears amenable to a wide range of kernel markers and phenotypes (for example, those found in Figure 3A). Additional phenotypes will require additional training data, but no technical limitations exist for the addition of kernel markers such as anthocyanin genes or kernel morphology markers like *bt1*. Agriculturally relevant aspects of maize ears, such as kernel size, row number, and other yield components, could also conceivably be measured with images generated by the ear scanner. The addition of calibration surfaces (such as a ruler or grid) to the scanner would likely improve the ability to estimate such ear or kernel dimensions.

Images of ears provide a convenient, long-lasting record of experiments, particularly if they are shared by researchers. These data can be used in a variety of ways, such as measuring patterns of kernel distribution, quantifying empty space on the ear, and recording other phenotypes such as abnormal or aborted kernels. For our experimental objectives, the system made it feasible to generate a nearly two-fold larger set of fluorescent kernel transmission data compared to our initial study (Warman et al. 2020). Manual quantification of the original dataset took approximately 50 hours. Automated quantification of the larger dataset using EarVision took less than four hours when run on multiple GPUs, representing an approximately 25-fold decrease in the time required to quantify the images. Not only did the larger dataset confirm the observations in that study, it also enabled the identification of a new male-specific gametophytic mutant, pointing toward a previously unknown function for a vacuolar processing enzyme. The maize ear scanning and EarVision system increases the scope of feasible experiments addressing maize reproductive biology and related agricultural traits by addressing a bottleneck in data acquisition and quantification, paving the way for high-throughput phenotyping in this area.

## Materials and Methods

### Building the maize ear scanner

The maize ear scanner was built from dimensional lumber and widely available parts. For detailed plans and three-dimensional models, see Supplemental File 1. The base of the scanner was built from a nominal 2×12 (38×286 mm) fir board, while the frame of the scanner was built from nominal 2×2 (38×38 mm) cedar boards. Boards were fastened together with screws. Strict adherence to materials and exact dimensions of the scanner frame is not necessary, as long as the scanner is structurally sound and large enough to accommodate ears of varying sizes.

A standard rotisserie motor (Minostar universal grill electric replacement rotisserie motor, 120 volt 4 watt), used to rotate the maize ear, was attached to the base of the scanner by way of a wood enclosure. A 5/16” (8 mm) steel rod was placed in the rotisserie motor to provide a point to fasten the lower portion of the ear. The top of the steel rod was ground to a flattened point with a bench grinder to allow it to be inserted into the pith at the center of the base of the ear.

The top of the ear was held in place with an adjustable assembly constructed from a nominal 2×4 board (38×89 mm) fastened to drawer slides (Liberty D80618C-ZP-W 18-inch ball bearing drawer slides) on either side of the scanner frame (Supplemental File 1). In the center of the 2×4, facing down towards the top of the ear, is a steel pin mounted on a pillow block bearing (Letool 12mm mounted housing self-aligning pillow flange block bearing). The steel pin (12mm) was sharpened to a point to penetrate the top of the ear as it is lowered, temporarily holding it in place while the ear is rotated during scanning. Because the pin can be moved up and down on the drawer slides, a variety of ear sizes can be accommodated in the scanner.

Ambient lighting was used for full spectrum visible light images. To capture GFP fluorescence, a blue light (Clare Chemical HL34T) was used to illuminate the ear. An orange filter (Neewer camera flash color gel kit) was placed in front of the camera lens to partially filter out non-GFP wavelengths.

### Ear scanning workflow

Preparation for the scanning process begins by trimming the top and bottom of the ear to expose the central pith. Before scanning, ear dimensions (length and diameter at widest point) were recorded. Following measurement, the bottom pin is inserted into the bottom of the ear, after which the pin with ear attached is placed in the rotisserie motor. The top of the ear is secured by lowering the top pin into the pith at the top of the ear.

Ear scanning was divided into two configurations. In the first configuration (MES v1.0), a camera capable of capturing videos (such as a cell phone or point-and-shoot digital camera, a Sony DSCWX220 was used in our version) was mounted on a tripod approximately 60 cm in front of the rotating ear. Videos were captured by the camera, then manually imported to a computer for processing and downstream analysis. In the second configuration (MES v2.0), a USB camera (ELP USBFHD06H-SFV) capable of capturing 1080p resolution video at 30 fps is directly controlled by a desktop computer (Dell 3630) running the Ubuntu Linux distribution (version 18.04.3). The camera was placed approximately 60 cm in front of the ear for video capture. Videos are previewed using the command line utility qv4l2 (V4L2 Test Bench, version 1.10.0), then captured using a custom FFmpeg command (ffmpeg -t 27 -f v4l2 -framerate 30 - video_size 1920×1080 -i/dev/video1/output.mov). The command captures the number of frames required for one complete rotation of the ear plus a small initial buffer. Videos are processed into flat images each night by running a custom script (see following section) with the Linux cron utility. After video processing, flat images are uploaded to Google cloud and local server space using the rclone (version 1.50.2) and rsync (version 3.1.2) utilities, respectively. A detailed protocol for scanning ears with the maize ear scanner using ears with GFP kernel markers and an ELP USBFHD06H-SFV USB camera can be found in Supplemental File 2.

### Creating flat images

Videos were processed to flat images. Frames were first extracted from videos to png formatted images using FFmpeg with default options (ffmpeg -i./”$file” -threads 4./maize_processing_folder/output_%04d.png). These images were then cropped to the central row of pixels using ImageMagick (mogrify -verbose -crop 1920×1+0+540 +repage./maize_processing_folder/*.png). The collection of single pixel row images was then appended in sequential order (convert -verbose -append +repage./maize_processing_folder/*.png./”$name.png”). Finally, the image was rotated and cropped (mogrify -rotate “180” +repage./”$name.png”; mogrify -crop 1920×746+0+40 +repage./”$name.png”). We chose the convention of a horizontal flattened image with the top of the ear to the right and the bottom of the ear to the left. Because the videos were captured vertically, a rotation was required after appending the individual frames. The vertical dimension of the final crop reflects the number of frames (746) required for one full rotation of the ear.

### Manually quantifying kernels using flat images

Kernels were quantified from ear projections using the Cell Counter plugin of the FIJI distribution of ImageJ (version 2.0.0) (Schindelin et al., 2012). Ears were assigned counter types to correspond to different kernel markers, after which kernels on ear images were located and annotated manually. The Cell Counter plugin exports results in an xml file, which contains the coordinates and marker type of every annotated kernel. This file can be processed to create a map of kernel locations on the ear. A detailed protocol describing the quantification process can be found in Supplemental File 3.

### Image segmentation and labeling by watershed transformation and k-means clustering

Two-dimensional projections of images containing GFP kernel markers were segmented using a watershed transformation implemented in the scikit-image Python library, version 0.16.1. The tutorials located at https://scikit-image.org/docs/stable/auto_examples/applications/plot_coins_segmentation.html and https://scikit-image.org/docs/dev/auto_examples/color_exposure/plot_regional_maxima.html were used as starting points. Images were first cropped by 15% along each side to remove distorted regions along the top and bottom of the ear. Next, regional maxima were isolated from the images using the scikit-image “reconstruction” function with the original image minus a fixed h-value of 0.3 as the seed image. The resulting h-dome regional maxima were further processed using the Sobel operator (scikit-image “sobel” function). Extreme high values of the resulting image’s histogram were used as seeds for the scikit-image “watershed” function. Finally, connections between adjacent kernel segments were reduced by morphological opening using the “binary_opening” scikit-image function.

Once segments identifying potential kernels were identified, segments were classified into either “fluorescent” or “non-fluorescent” categories. First, segment centers were identified using the “center_of_mass” function from the SciPy Multi-dimensional image processing package (version 1.4.1). Mean intensity in red, green, and blue channels was then calculated for each segment. Segments were divided into two clusters by channel intensity by k-means clustering using the “kmeans” function from the scikit-learn library (version 0.22.2). Clusters were collectively identified as the “fluorescent” or “non-fluorescent” cluster based on their relative mean segment intensity in the green channel. Fluorescent, non-fluorescent, and percent fluorescent metrics were calculated using this method for 320 images in the test dataset described in the following section. Adjusted R^2^ values were calculated using a linear regression for each metric.

### Training, validation, and test dataset generation

Training and validation datasets were generated from 300 scanned ear images from the 2018 and 2019 field seasons (150 images each season), using a 0.7 training to validation ratio. Lines contained a selection of single mutations from the Dooner/Du collection of *Ds-GFP*-tagged transposable element insertions (Li et al., 2013). Kernels were manually annotated with bounding boxes and classes (fluorescent or non-fluorescent) using LabelImg (https://github.com/tzutalin/labelImg).

A test dataset was generated using 320 scanned images of *Ds-GFP-tagged* ears from the 2018 and 2019 field seasons (160 images each season). A Sony DSCWX220 camera was used to capture images in 2018 and an ELP USBFHD06H-SFV was used to capture images in 2019. Images used for training and validation of the model were excluded from the test dataset. Total fluorescent and non-fluorescent kernels were quantified using ImageJ as previously described (see section “Manually quantifying kernels using flat images”).

### Deep learning model selection and configuration

The deep learning pipeline used the Faster R-CNN with Inception Resnet v2 with Atrous convolutions model, implemented in the Tensorflow Object Detection API. A repository containing the code used to train the model and run inference, titled EarVision, is linked below. To preserve GPU memory, images were resized to maximum dimensions of 600×1024 pixels. Training data was split into training and validation sets with a ratio of training to validation of 0.7. First stage RPN anchor proposals were limited to 3000, with 8 aspect ratios at each anchor point. Max total detections were set at 2000. Data augmentations were limited to a random horizontal flip. For full configuration file, see EarVision repository.

Models were created using two approaches. In the first, a single model was trained with data from combined 2018 and 2019 field seasons. Separate cameras were used for each season (see description in the previous section). This model was trained for 940 epochs on an Nvidia V100 GPU, a process that took approximately 74 hours. The training length was determined by optimizing mAP (mean average precision) at 0.5 IOU (intersection over union). This parameter measures the average precision (true positives divided by the sum of true positives and false positives) over a range of recall values (true positives divided by the sum of true positives and false negatives). This metric summarizes the model’s performance at correctly identifying bounding boxes and classes while minimizing false positives. At epoch 940, the model’s mAP at 0.5 IOU was 0.790, with an average recall of 0.622.

In the second approach, two models were trained independently on 2018 and 2019 images. These models were trained for 2226 and 2468 epochs, respectively, for approximately 45 minutes on an Nvidia V100 GPU. Training lengths were optimized as with the single model described previously. The 2018 and 2019 mAP at 0.5 IOU were approximately 0.843 and 0.867, respectively, with average recalls of 0.601 and 0.693.

### Image subdivision and bounding box confidence scores

Before training, images and annotations were divided into three sub-images using a custom script (see below). For inference, input images were likewise divided into three subimages. In both cases, images were divided along the horizontal axis (Figure 4B). Overlapping regions of 100 pixels in width (all pixel measurements based on non-scaled input images, generally 1920×746 pixels) were included in the left and right sub-images. Because empty margins on the left and right of the original image generally led to the center sub-image having the largest number of kernels, only the left and right sub-images included 100 pixels overlapping regions.

Inference was first run on each sub-image individually. Next, bounding boxes within 40 pixels of subdivision borders were removed. This process removed partial bounding boxes of kernels located along the dividing lines between images. Because of image overlap, these kernels were still marked by complete bounding boxes after partial bounding boxes were removed. After this step, bounding boxes and annotations from the three sub-images were combined. Redundant bounding boxes in overlapping regions were removed by non-maximum suppression using the Tensorflow function “non_max_suppression”. Non-maximum suppression calculates the intersection over union (IOU) value for all bounding box pairs. For pairs that exceed a defined intersection over union (IOU) value, in our case 0.5, the bounding box with the lowest confidence score is removed. Inference for each input image took approximately one minute on a Nvidia M10 GPU, with individual models performing slightly faster than the single model.

Optimal confidence score thresholds for final bounding box outputs were determined empirically by maximizing the R-squared value for total fluorescent and non-fluorescent kernel count across the test image dataset. R-squared values for confidence thresholds ranging from 0-1 in 0.01 increments were calculated for both fluorescent and non-fluorescent total kernel counts by comparing model predictions and manually validated data (Supplemental Figure 5). A single confidence threshold of 0.12 was chosen for the combined 2018/2019 model to maximize the combined R-squared value in both classes (Supplemental Figure 5A). Confidence thresholds of 0.08 and 0.12 were chosen for 2018 and 2019 individual models (Supplemental Figure 5B-C).

### Statistical methods for deep learning model application to test dataset

Manually counted kernel totals were compared with deep learning model predictions for the 320 test images by fitting a linear regression using the “lm” function in R. Adjusted R^2^ values were calculated for fluorescent and non-fluorescent kernels, as well as for percent fluorescent kernel transmission. Mean absolute deviations were calculated for fluorescent and non-fluorescent total kernel counts, and percent fluorescent kernel transmission. Analysis was carried out using both a single model trained on 2018 and 2019 images, as well as individual models trained on each year alone.

### Experimental design and statistical methods for deep learning model application to 2018 and 2019 field trials

Inference was run on 983 scanned images from the 2018 (369 images) and 2019 (614 images) field seasons (Supplemental Table 1). Scanned ear images were the result of reciprocal outcrosses of heterozygous plants carrying GFP-tagged *Ds* insertion alleles in a variety of genes highly expressed across maize gametophyte development. For detailed experimental description, see (Warman et al., 2020). In brief, alleles were chosen from highly expressed genes (top 20% by FPKM) in three categories: Vegetative Cell, Sperm Cell, and Seedling as a sporophytic control. A total of 56 alleles were quantified in (Warman et al., 2020), of which 48 displayed fluorescent seed markers and were analyzed in this study. Eight alleles were associated with anthocyanin seed markers, and were thus not included in this analysis. Ear images from the 2019 field season contained additional crosses from the alleles present in the 2018 field season, plus 12 additional alleles (summarized in Table 2), for a total of 60 alleles.

After model inference, total fluorescent and non-fluorescent seed counts were analyzed using a generalized linear model with a logit link function for binomial counts and a quasibinomial family to correct for overdispersion between parent lines. Significant differences from expected 50% inheritance were assessed with a quasi-likelihood test with p-values corrected for multiple testing using the Benjamini-Hochberg procedure to control the false discovery rate at 0.05. Significant differences from 50% inheritance were defined with an adjusted p-value < 0.05. Separate generalized linear models were carried out for each year and cross category (female, Seedling male, Vegetative Cell male, Sperm Cell male). A combined generalized linear model with all 2018 manual counts and all 2018 computer vision predictions was also created in order to determine the significance of manual counts versus computer vision predictions as a factor.

## Supporting information

Rotational ear scanner construction plans.

Scanning fluorescent ears with the rotational ears scanner protocol

Quantifying kernels in flat images using ImageJ protocol

Final EarVision kernel count predictions for images of crosses generated in two field seasons

## Supplemental Data

**Supplemental Figures.** Supplemental Figures and Figure Legends.

**Supplemental File 1.** Rotational ear scanner construction plans.

**Supplemental File 2.** Scanning fluorescent ears with the rotational ears scanner protocol.

**Supplemental File 3.** Quantifying kernels in flat images using ImageJ protocol.

**Supplemental Table 1.** Final EarVision kernel count predictions for images of crosses generated in two field seasons, as shown in Figure 6.

## Software and image data availability

### Ear video processing to flat images

This script processes videos from the MES into flat ear projection images. (https://github.com/fowler-lab-osu/make_flat_images_from_videos)

### Traditional computer vision methods

This repository contains code used to segment kernels from images, described in the section “A traditional computer vision approach for automated discrimination of fluorescent and wild-type kernels.” (https://github.com/fowler-lab-osu/traditional_cv_kernel_counter)

### Statistical methods

This repository contains statistical methods used to analyze data and generate figure plots in R. (https://github.com/fowler-lab-osu/maize_ear_scanner_and_computer_vision_statistics)

### EarVision

These repositories contain the EarVision computer vision pipeline for kernel identification (https://github.com/fowler-lab-osu/EarVision and https://github.com/fowler-lab-osu/EarVision_TensorFlow_Object_Detection_API)

### Training and validation images

This set of images includes those used for training and validation of the EarVision model, with a total of 300 images and associated kernel annotations in Pascal VOC format. (http://files.cgrb.oregonstate.edu/Fowler_Lab/EarVision_maize_kernel_image_data/training_and_validation_images/)

### Testing images

This set of images includes those used for testing the EarVision model, with a total of 320 images. (http://files.cgrb.oregonstate.edu/Fowler_Lab/EarVision_maize_kernel_image_data/testing_images/)

### Example large scale application images

This set of images includes those used in the section, “Application of deep learning models to a large ear projection dataset,” with a total of 983 images. (http://files.cgrb.oregonstate.edu/Fowler_Lab/EarVision_maize_kernel_image_data/example_large_scale_application_images/)

## Acknowledgements

We thank O. Childress, H. Fowler, B. Galardi, B. Hamilton, R. Hartman, K. Kress, C. Lambert, and M. Wesel for their annotation and kernel counting assistance. In addition, we thank O. Childress for protocol feedback and E. Vischulis for technical assistance. Thank you to Jeff H. Chang for reading the manuscript and providing useful comments. This work was funded by National Science Foundation grants IOS-1340050 and MCB-1832186 (to JEF) and IOS-1340112 (PJ), as well as the Department of Botany & Plant Pathology at Oregon State University. Computing resources were provided by the Center for Genome Research and Biocomputing (CGRB) at Oregon State University. Oregon State University has filed a provisional patent application (pending, # 62/989,403) in support of commercialization of this technology.

## Literature Cited

Abadi M, Agarwal A, Barham P, Brevdo E, Chen Z, Citro C, Corrado GS, Davis A, Dean J, Devin M, et al (2016a) TensorFlow: Large-Scale Machine Learning on Heterogeneous Distributed Systems. arXiv [cs.DC]

Abadi M, Barham P, Chen J, Chen Z, Davis A, Dean J, Devin M, Ghemawat S, Irving G, Isard M, et al (2016b) TensorFlow: A System for Large-Scale Machine Learning. 12th USENIX Symposium on Operating Systems Design and Implementation (OSDI 16). pp 265–283

Angermueller C, Pärnamaa T, Parts L, Stegle O (2016) Deep learning for computational biology. Mol Syst Biol 12: 878

Arthur KM, Vejlupkova Z, Meeley RB, Fowler JE (2003) Maize ROP2 GTPase provides a competitive advantage to the male gametophyte. Genetics 165: 2137–2151

Bai F, Daliberti M, Bagadion A, Xu M, Li Y, Baier J, Tseung C-W, Evans MMS, Settles AM (2016) Parent-of-Origin-Effect rough endosperm Mutants in Maize. Genetics 204: 221–231

Chaivivatrakul S, Tang L, Dailey MN, Nakarmi AD (2014) Automatic morphological trait characterization for corn plants via 3D holographic reconstruction. Comput Electron Agric 109: 109–123

Ching T, Himmelstein DS, Beaulieu-Jones BK, Kalinin AA, Do BT, Way GP, Ferrero E, Agapow P-M, Zietz M, Hoffman MM, et al (2018) Opportunities and obstacles for deep learning in biology and medicine. J R Soc Interface. doi: 10.1098/rsif.2017.0387

Choudhury SD, Stoerger V, Samal A, Schnable JC, Liang Z, Yu J-G (2016) Automated vegetative stage phenotyping analysis of maize plants using visible light images. KDD workshop on data science for food, energy and water, San Francisco, California, USA

Clark RT, Famoso AN, Zhao K, Shaff JE, Craft EJ, Bustamante CD, McCouch SR, Aneshansley DJ, Kochian LV (2013) High-throughput two-dimensional root system phenotyping platform facilitates genetic analysis of root growth and development. Plant Cell Environ 36: 454–466

Dobos O, Horvath P, Nagy F, Danka T, Viczián A (2019) A Deep Learning-Based Approach for High-Throughput Hypocotyl Phenotyping. Plant Physiol 181: 1415–1424

Fahlgren N, Gehan MA, Baxter I (2015) Lights, camera, action: high-throughput plant phenotyping is ready for a close-up. Curr Opin Plant Biol 24: 93–99

Feldmann M, Tabb A, Knapp SJ (2019) Cost-effective, high-throughput 3D reconstruction method for fruit phenotyping. Computer Vision Problems in Plant Phenotyping (CVPPP) 1

Gibbs JA, Burgess AJ, Pound MP, Pridmore TP, Murchie EH (2019) Recovering Wind-Induced Plant Motion in Dense Field Environments via Deep Learning and Multiple Object Tracking. Plant Physiol 181: 28–42

Huang J, Rathod V, Sun C, Zhu M, Korattikara A, Fathi A, Fischer I, Wojna Z, Song Y, Guadarrama S, et al (2016) Speed/accuracy trade-offs for modern convolutional object detectors. arXiv [cs.CV]

Huang JT, Wang Q, Park W, Feng Y, Kumar D, Meeley R, Dooner HK (2017) Competitive Ability of Maize Pollen Grains Requires Paralogous Serine Threonine Protein Kinases STK1 and STK2. Genetics 207: 1361–1370

Jiang N, Floro E, Bray AL, Laws B, Duncan KE, Topp CN (2019) Three-Dimensional TimeLapse Analysis Reveals Multiscale Relationships in Maize Root Systems with Contrasting Architectures. Plant Cell 31: 1708–1722

Junker A, Muraya MM, Weigelt-Fischer K, Arana-Ceballos F, Klukas C, Melchinger AE, Meyer RC, Riewe D, Altmann T (2014) Optimizing experimental procedures for quantitative evaluation of crop plant performance in high throughput phenotyping systems. Front Plant Sci 5: 770

Liang X, Wang K, Huang C, Zhang X, Yan J, Yang W (2016) A high-throughput maize kernel traits scorer based on line-scan imaging. Measurement 90: 453–460

Li Y, Segal G, Wang Q, Dooner HK (2013) Gene Tagging with Engineered Ds Elements in Maize. In T Peterson, ed, Plant Transposable Elements: Methods and Protocols. Humana Press, Totowa, NJ, pp 83–99

Mahlein A-K (2016) Plant Disease Detection by Imaging Sensors -Parallels and Specific Demands for Precision Agriculture and Plant Phenotyping. Plant Dis 100: 241–251

Makanza R, Zaman-Allah M, Cairns JE, Eyre J, Burgueño J, Pacheco Á, Diepenbrock C, Magorokosho C, Tarekegne A, Olsen M, et al (2018) High-throughput method for ear phenotyping and kernel weight estimation in maize using ear digital imaging. Plant Methods 14: 49

Miller ND, Haase NJ, Lee J, Kaeppler SM, de Leon N, Spalding EP (2017) A robust, high-throughput method for computing maize ear, cob, and kernel attributes automatically from images. Plant J 89: 169–178

Mohanty SP, Hughes DP, Salathé M (2016) Using Deep Learning for Image-Based Plant Disease Detection. Front Plant Sci 7: 1419

Neuffer MG, Coe EH, Wessler SR (1997) Mutants of maize. Cold Spring Harbor Laboratory Press

Phillips AR, Evans MMS (2011) Analysis of stunter1, a maize mutant with reduced gametophyte size and maternal effects on seed development. Genetics 187: 1085–1097

Rawat W, Wang Z (2017) Deep Convolutional Neural Networks for Image Classification: A Comprehensive Review. Neural Comput 29: 2352–2449

Redmon J, Farhadi A (2018) YOLOv3: An Incremental Improvement. arXiv [cs.CV]

Ren S, He K, Girshick R, Sun J (2015) Faster R-CNN: Towards Real-Time Object Detection with Region Proposal Networks. In C Cortes, ND Lawrence, DD Lee, M Sugiyama, R Garnett, eds, Advances in Neural Information Processing Systems 28. Curran Associates, Inc., pp 91–99

Schindelin J, Arganda-Carreras I, Frise E, Kaynig V, Longair M, Pietzsch T, Preibisch S, Rueden C, Saalfeld S, Schmid B, et al (2012) Fiji: an open-source platform for biological-image analysis. Nat Methods 9: 676–682

Szegedy C, Ioffe S, Vanhoucke V, Alemi A (2016) Inception-v4, Inception-ResNet and the Impact of Residual Connections on Learning. arXiv [cs.CV]

Tardieu F, Cabrera-Bosquet L, Pridmore T, Bennett M (2017) Plant Phenomics, From Sensors to Knowledge. Curr Biol 27: R770–R783

Ubbens JR, Stavness I (2017) Deep Plant Phenomics: A Deep Learning Platform for Complex Plant Phenotyping Tasks. Front Plant Sci 8: 1190

Warman C, Panda K, Vejlupkova Z, Hokin S, Unger-Wallace E, Cole RA, Chettoor AM, Jiang D, Vollbrecht E, Evans MMS, et al (2020) High expression in maize pollen correlates with genetic contributions to pollen fitness as well as with coordinated transcription from neighboring transposable elements. PLoS Genet 16: e1008462

Wen W, Guo X, Lu X, Wang Y, Yu Z (2019) Multi-scale 3D Data Acquisition of Maize. Computer and Computing Technologies in Agriculture XI. Springer International Publishing, pp 108xyo115

Yamada K, Shimada T, Nishimura M, Hara-Nishimura I (2005) A VPE family supporting various vacuolar functions in plants. Physiol Plant 123: 369–375

Zhang X, Huang C, Wu D, Qiao F, Li W, Duan L, Wang K, Xiao Y, Chen G, Liu Q, et al (2017) High-Throughput Phenotyping and QTL Mapping Reveals the Genetic Architecture of Maize Plant Growth. Plant Physiol 173: 1554–1564

