## Supplemental Figures and Figure Legends for "A cost-effective maize ear phenotyping platform enables rapid categorization and quantification of kernels"

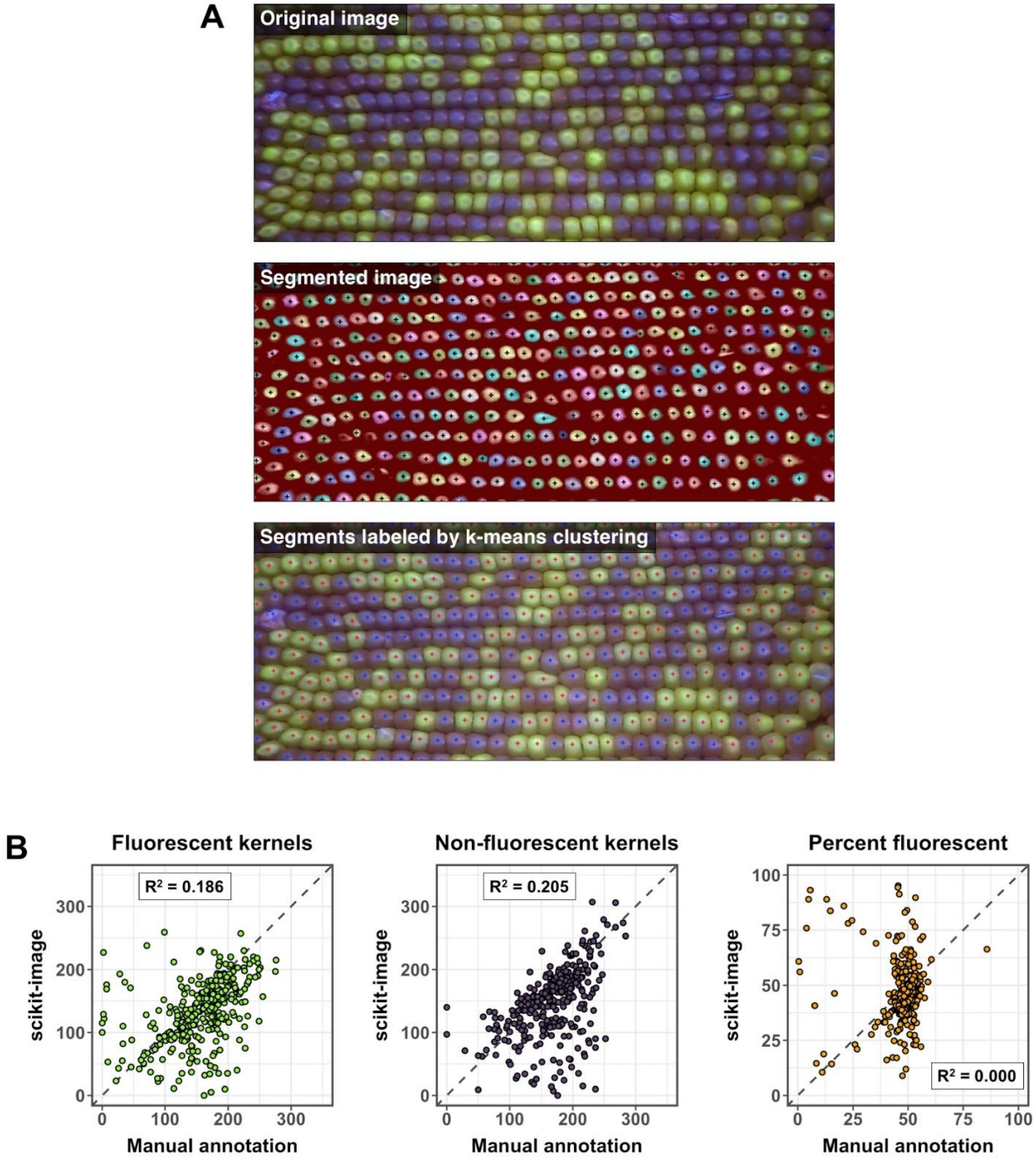


**Supplemental Figure 1.** A traditional computer vision approach generalized poorly for automated kernel detection.

**(A)** Top panel: Two-dimension maize ear projection showing fluorescent and non-fluorescent kernels (GFP kernel marker). Middle panel: Watershed transformation followed by

morphological opening segments individual kernels. Bottom panel: Mean intensity of RGB color values within each segment are clustered by k-means clustering into two groups,  fluorescent and non-fluorescent.
**(B)** Watershed computer vision approach applied to combined 2018 + 2019 ear image test sets. Diagonal dashed line indicates equal manual counts and computer vision predictions.

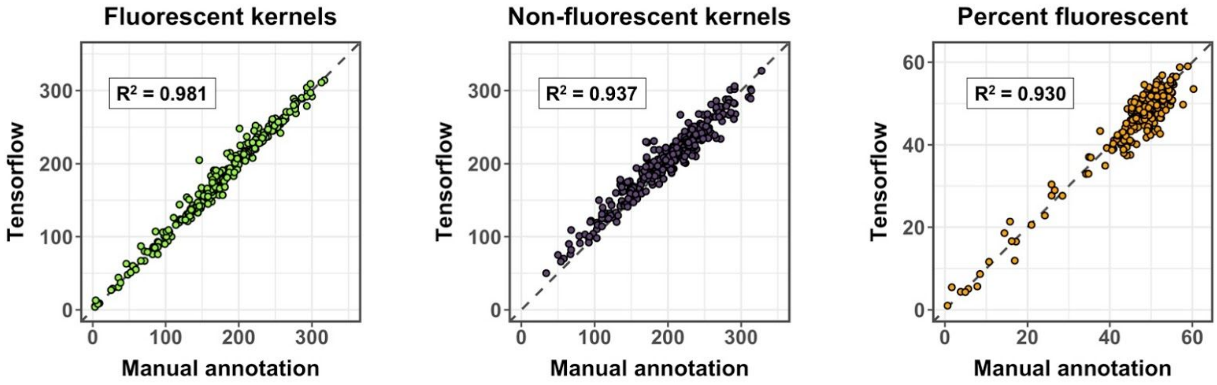


**Supplemental Figure 2.** A single deep learning model trained on images from two different cameras was able to detect kernels with moderate success.

Plots comparing model predictions and manual counts in the ear image test dataset. Model predictions were made using a model trained on images from two different cameras. Dashed diagonal lines indicate equal manual counts and model predictions.

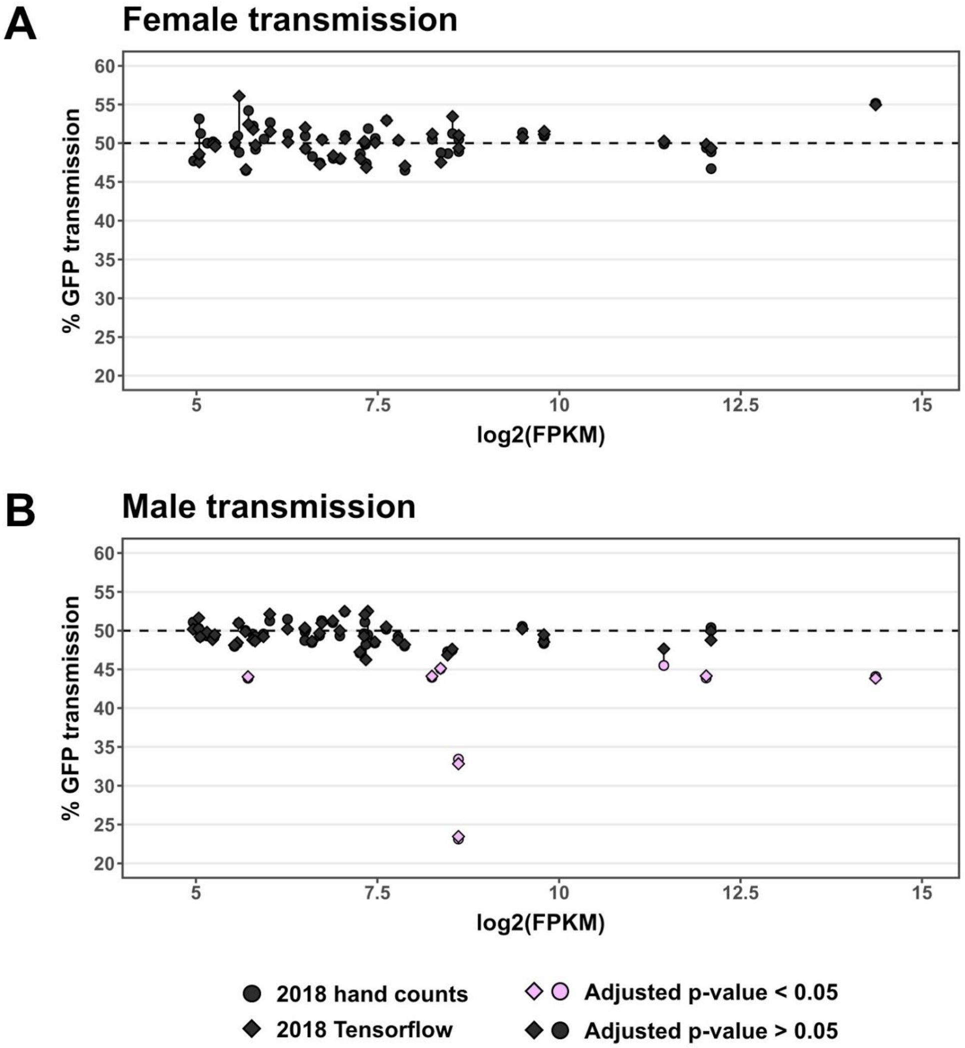


**Supplemental Figure 3.** Comparing 2018 hand counts to 2018 Tensorflow model predictions.

**(A)** GFP transmission in female crosses. No model-predicted transmission rates were significantly different from Mendelian inheritance, in agreement with hand counts. **(B)** GFP transmission in male crosses. While 7 out of 8 alleles with significant transmission defects by hand counts were identified by model predictions, 1 allele that was determined to be significant by hand counts was predicted to be non-significantly different than Mendelian by our model.

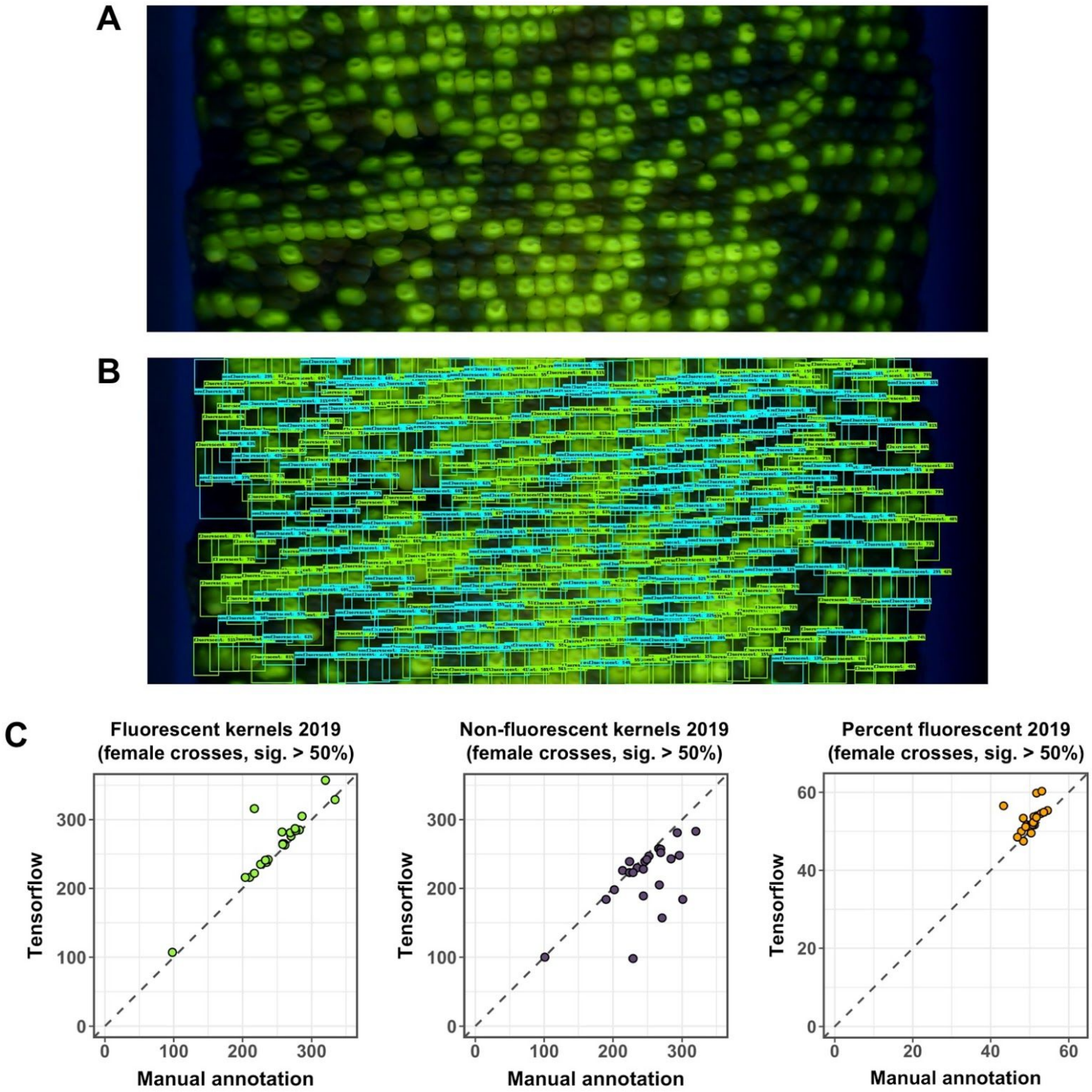


**Supplemental Figure 4.** Underexposure of a subset of 2019 images led to false positives for female transmission

**(A)** Original image for a female cross from *Ds-GFP* allele *tdsgR84A12*, an insertion in maize gene Zm00001d005781.

**(B)** Model predicted bounding boxes.

**(C)** Model predictions for female crosses in the four alleles that had significantly over 50% inheritance, compared to manual annotations. Two images had substantial overestimations of GFP kernel totals (left), likely the result of strong fluorescence illuminating adjacent non-fluorescent kernels in these images. Underestimations of non-fluorescent kernels were widespread (middle). Overestimation of percent fluorescent kernels was present in a small subset of female images (right).

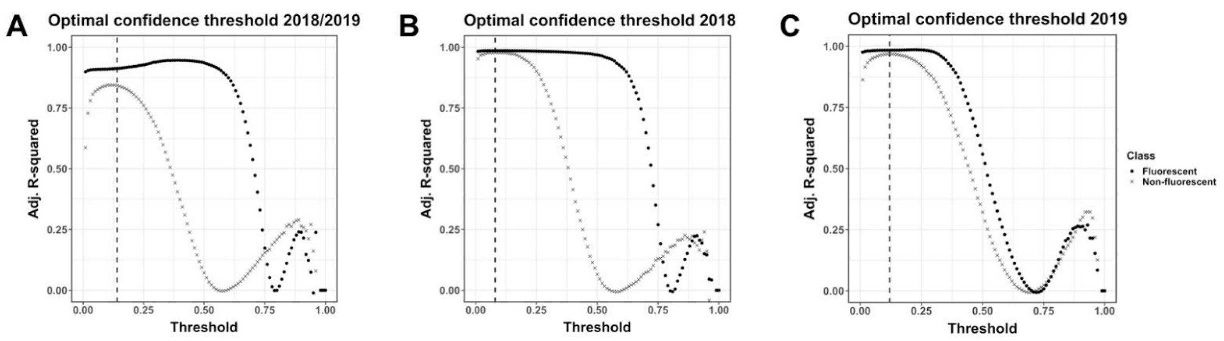


**Supplemental Figure 5.** Empirically determining optimal bounding box output confidence thresholds.

**(A)** Optimal confidence thresholds were predicted from test image sets for the combined 2018/2019 model. $R^2$ values were calculated for fluorescent and non-fluorescent classes by comparing total predicted kernels to ground truth data across confidence thresholds from 0 to 1 in 0.01 intervals. The highest combined $R^2$ value was found at a confidence threshold of 0.12, marked with a vertical dashed line. **(B)** Optimal confidence thresholds for 2018 model. The highest combined $R^2$ value was found at a confidence threshold of 0.08 (vertical dashed line). **(C)** Optimal confidence thresholds for 2019 model. The highest combined $R^2$ value was found at a confidence threshold of 0.12 (vertical dashed line). Confidence thresholds from A-C were used to output kernel counts and bounding boxes in the final models.
