## Supplementary figures and images for "A cost-effective maize ear phenotyping platform enables rapid categorization and quantification of kernels"

### image_of_scanner.png

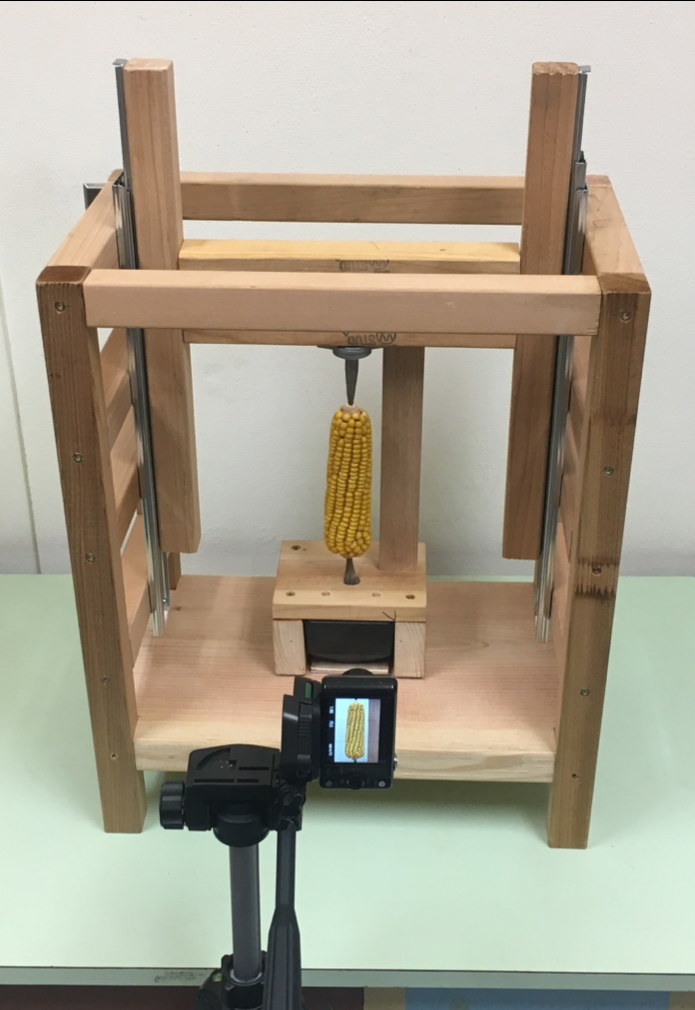

### scanner_plans_front_view.pdf

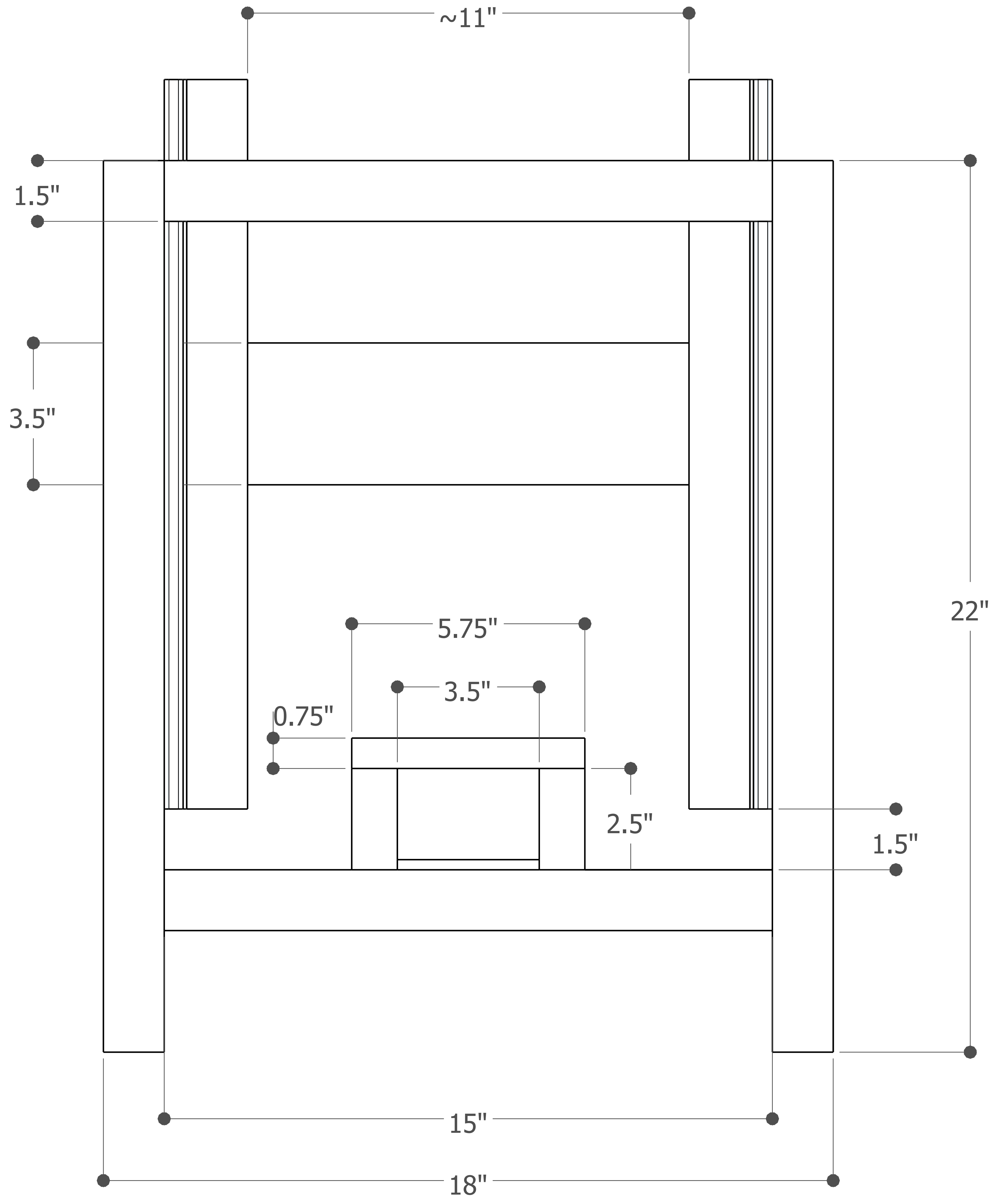

### scanner_plans_no_dimensions_closed.pdf

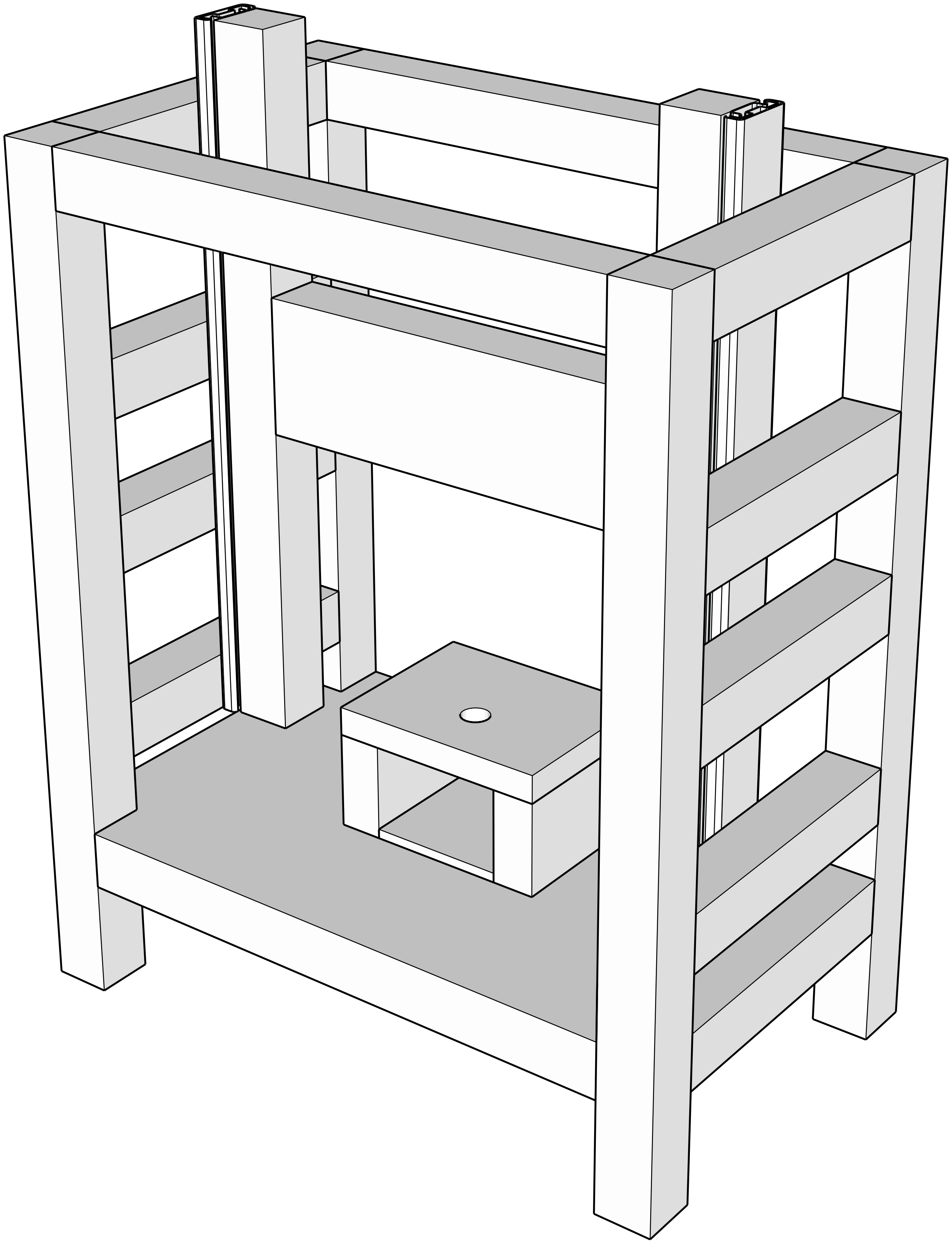

### scanner_plans_no_dimensions_open.pdf

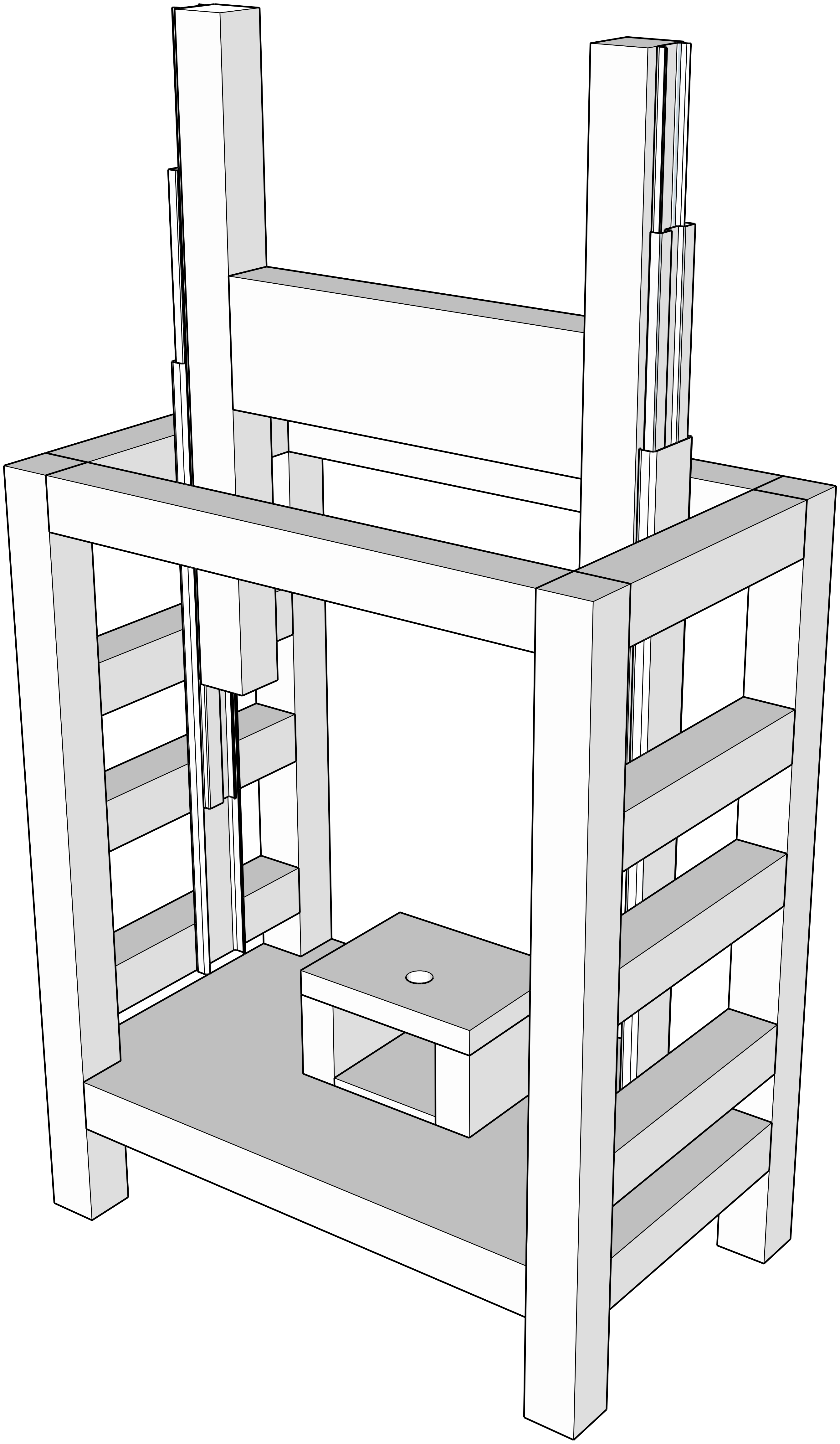

### scanner_plans_side_view.pdf

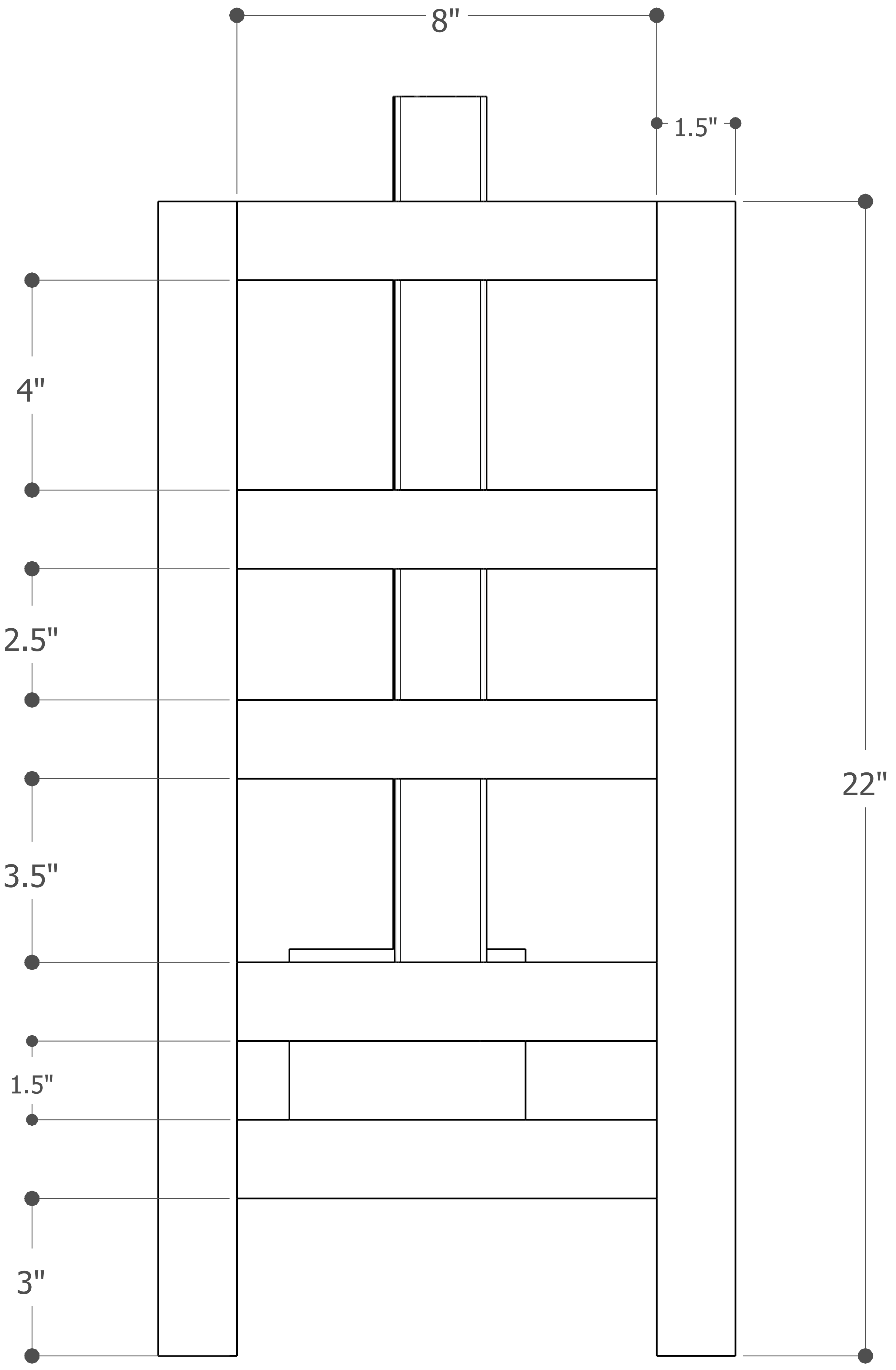

### scanner_plans_top_view.pdf

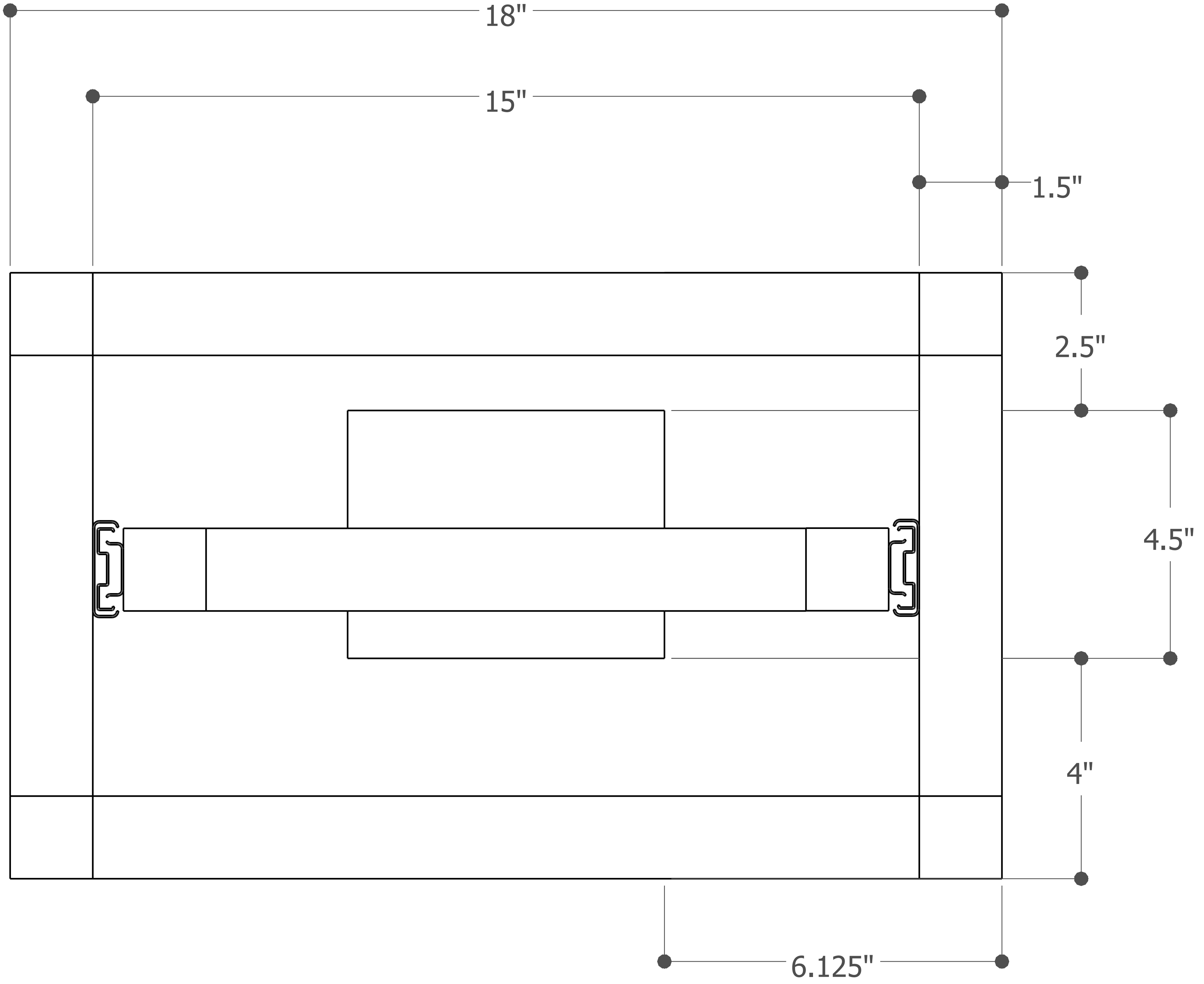

### scanner_plans_with_dimensions.pdf

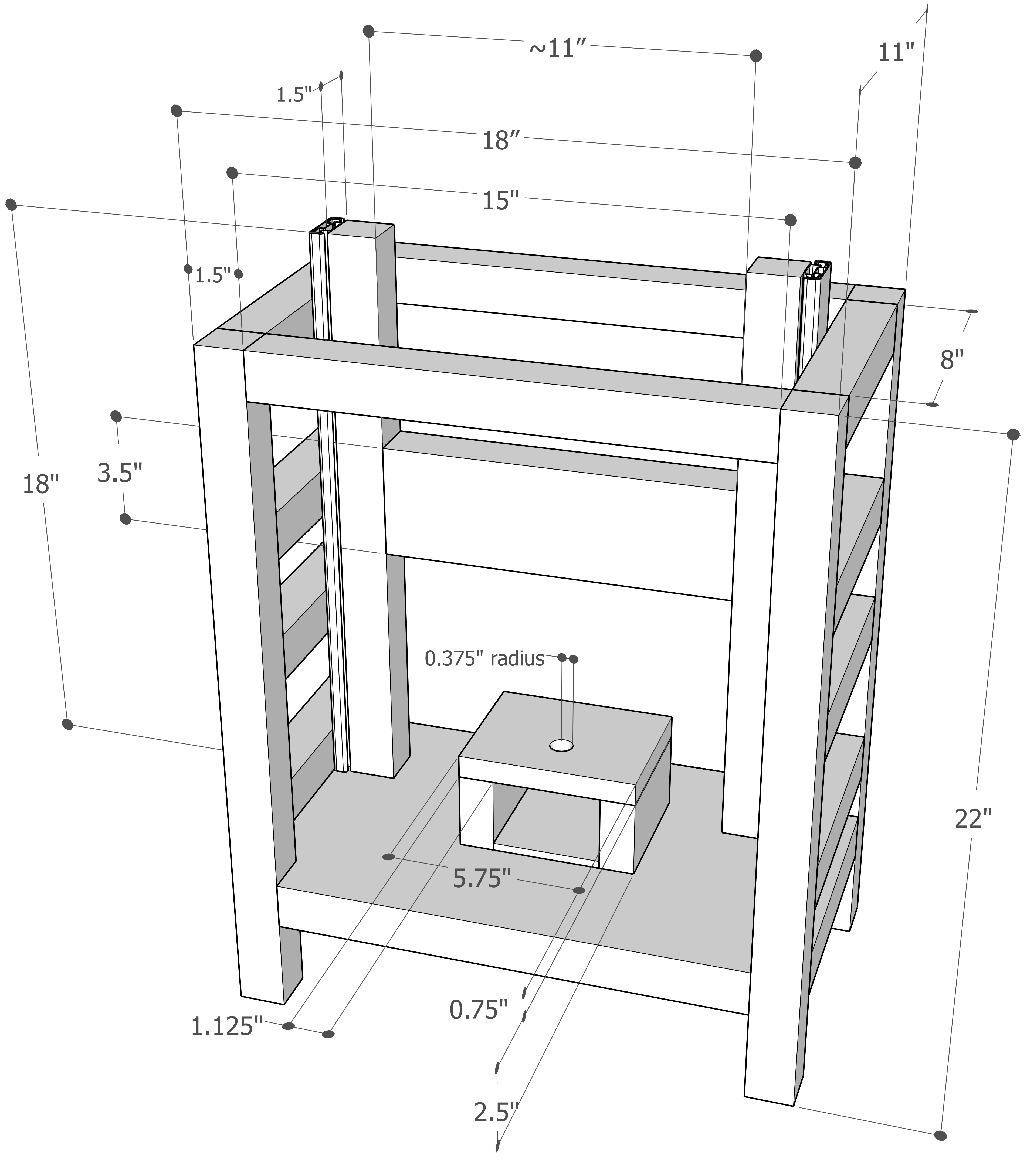
