## Supplementary material for "A cost-effective maize ear phenotyping platform enables rapid categorization and quantification of kernels": Scanning fluorescent ears with the rotational ears scanner protocol

### Scanning fluorescent ears with the maize ear scanner

*This protocol describes the process of scanning maize ears with green fluorescent kernels (dsGFP). A video is created of the rotating ear, which is then flattened into an image containing the entire surface of the ear. This version uses Ubuntu Linux, qv4l2, ffmpeg, and a USB camera (ELP USBFHD06H-SFV) to capture the videos.*

1. Cut excess material off the top and bottom of the ear. The pith in the center of the ear should be exposed on both the top and bottom of the ear. Remove silks.

**Before:**

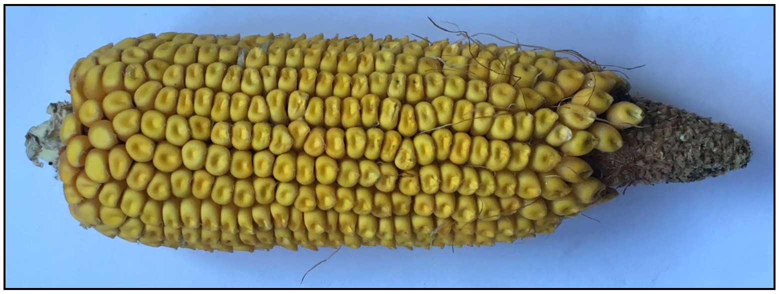

**After:**

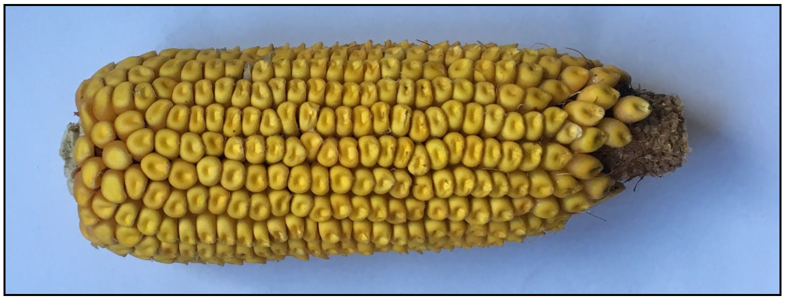

2. Insert bottom skewer into the pith on the base of the ear as shown, attempt to insert it as centered as possible. If you need to push hard to get it into the ear, use gloves.

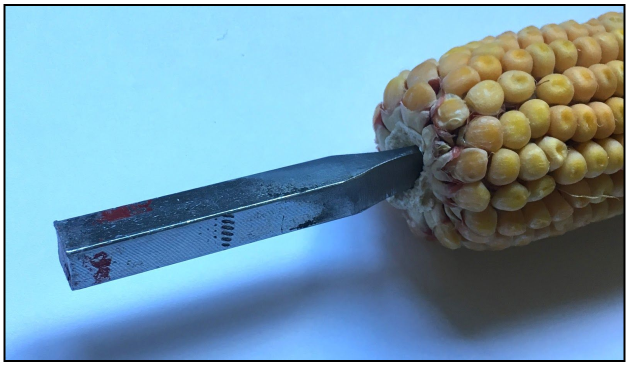

6. Raise top skewer and place bottom skewer in rotisserie.
7. Lower top skewer onto center of top of ear. Use caution when kernels go all the way to the top of the ear; it's possible to push too hard on the top skewer and forcibly dislodge several seeds. If kernels go all the way to the top of an ear, the top 10-20 kernels can be removed to make room for the top skewer. After removing the top kernels, trim the top of the ear to make a good position for the skewer.
8. Turn off the lights in the scanning room.
9. Adjust the blue light so that it shines perpendicular to the axis of ear rotation from the right side. The light should be vertically centered at the halfway point of the ear. In addition, place the light as close to the camera as possible, without the light obscuring any of the kernels.

**Optimal light placement:**

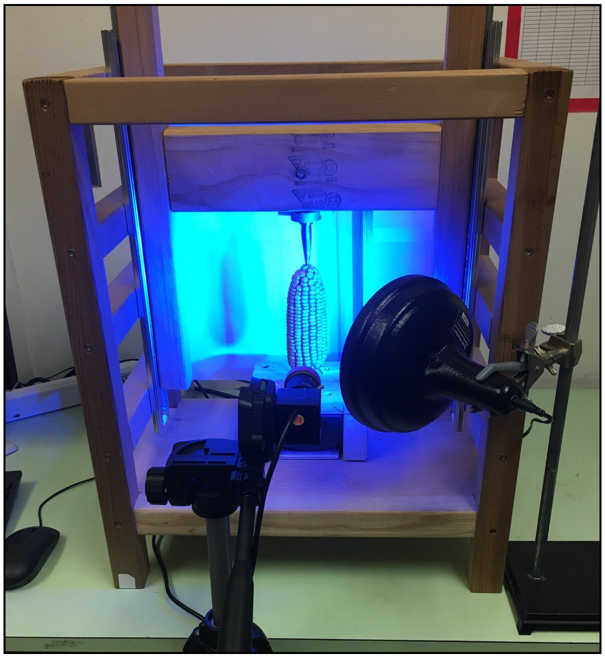

10. Begin previewing the camera image using the command line application qv4l2 (V4L2 Test Bench).
11. Ensure that the orange filter has been placed in front of the camera lense, and that glare from the light is minimized.
12. Ensure that the ear is centered in the preview window. The top and bottom of the ear should be close to the left and right side of the frame. Most importantly, the lengthwise center of the ear should be in the exact middle of the preview window.
13. Adjust exposure in the "Camera Controls" tab in the preview (note: if the video camera is currently running, close the video window to access the camera controls). For auto exposure, select "Aperture Priority Mode" in the Exposure, Auto dropdown menu. For manual exposure, select "Manual Mode." To change exposure in manual mode, type a number into the "Exposure (Absolute)" box. Between 300 and 600 seem to work well, but values will vary based on the ear and lighting conditions. Try to avoid exposures above 600 as the quality starts to degrade. For GFP videos, exposure should be long enough to highlight non-fluorescent seeds, but not so long that fluorescent seeds are overexposed (see examples below). For full spectrum videos, make sure the color seeds are clearly visible.

**Overexposed:**

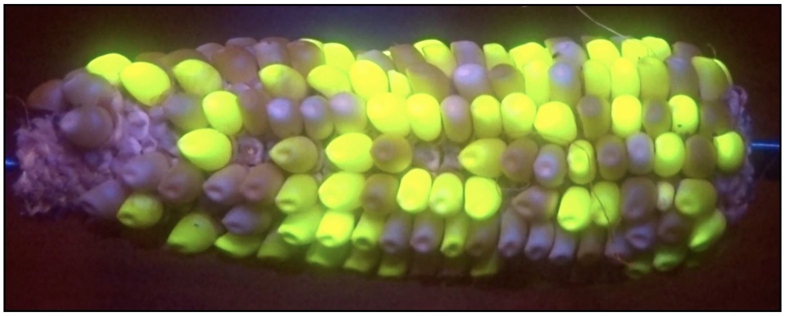

**Correct exposure:**

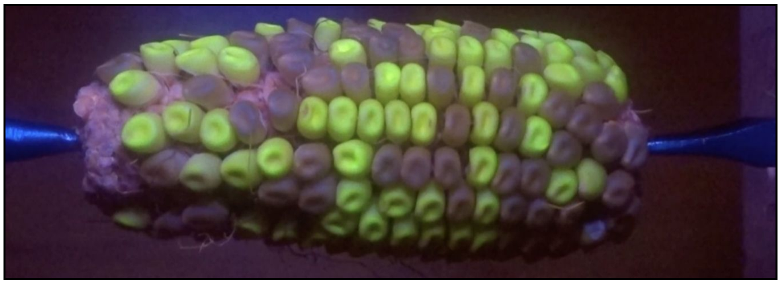

14. To capture video, close all preview windows and run the following ffmpeg command: ffmpeg -t 27 -f v4l2 -framerate 30 -video_size 1920x1080 -i /dev/video1 /output.mov (note: this command has been configured to capture the number of frames in one full rotation of our rotisserie motor, plus a small buffer. Different motor speeds will require modification of the -t parameter).
15. Remove ear from scanner, repeat for all ears.
